# Regulation of Renal Transporters by Pro-inflammatory Cytokines in Human Proximal Tubular Epithelial Cells: Identification of the Perpetrator and Mechanisms

**DOI:** 10.1101/2025.11.25.690608

**Authors:** Yik Pui Tsang, Kai Wang, Edward J. Kelly, Qingcheng Mao, Jashvant D. Unadkat

**Affiliations:** Department of Pharmaceutics, School of Pharmacy, University of Washington, Seattle, WA 98195, U.S.A; Kidney Research Institute, University of Washington, Seattle, Washington 98195, U.S.A

**Keywords:** transcription regulation, proximal tubule, pharmacokinetics, inflammation, gene expression, cytokines

## Abstract

**Introduction:** Infection and inflammation elevate circulating pro-inflammatory cytokines that can affect renal drug clearance. Accordingly, we sought to (i) quantify the extent of modulation of renal drug-metabolizing enzymes and transporters (DMETs) by cytokines and (ii) identify the mechanism(s) underlying these effects.

**Methods:** Fresh primary human proximal tubular epithelial cells (PTECs) were cultured on extracellular matrix-coated Transwells. PTECs were exposed every 24 h, for 48 h, to IL-6, IL-1β, TNF-α, IFN-γ, IL-4, or IL-10 (0.1 or 1 ng/mL), individually or as a cocktail. mRNA expression of 25 renal DMETs was quantified by RT-qPCR. Individual activity of OAT1–4, OCT2, and OCTN1 was measured. To determine mechanisms of these effects, selective MAPK/NF-κB inhibitors (ERK [PD98059], p38^MAPK^ [SB203580], JNK [SP600125], and NF-κB [PDTC]), individually or as a cocktail, were used. IL-6, soluble IL-6 receptor (sIL-6Rα), and IL-6 + sIL-6Rα were used to probe endogenous/exogenous IL-6 classic versus trans-signaling.

**Results:** IL-1β was the predominant modulator, downregulating mRNA expression of OAT1–3, OCT2, OAT4, MATE2-K, MRP2, and OATP4C1, and upregulating mRNA expression of OCTN1 and MRP3. TNF-α downregulated OAT1–3 mRNA expression to an extent similar to IL-1β, but did not affect other transporters. Activity changes for the major uptake transporters mirrored mRNA directionality. MAPK/NF-κB blockade by the inhibitor cocktail reduced IL-6 secretion while completely reversing the IL-1β-driven downregulation of OAT1–3 mRNA. JNK inhibition alone restored OAT1/3 mRNA. Inhibition of p38^MAPK^ blunted OAT2 mRNA downregulation. OCTN1 mRNA induction required NF-κB. Downregulation of OAT4/OCT2 mRNA was largely MAPK/NF-κB-independent. IL-6 alone, sIL-6Rα alone, or IL-6 + sIL-6Rα did not reproduce IL-1β-driven changes in transporter mRNA.

**Conclusions:** IL-1β is the principal driver of cytokine-mediated regulation of human renal transporters in PTECs via JNK/p38^MAPK^/NF-κB nodes. These mechanistic, exposure-verified data provide inputs for physiologically based pharmacokinetic predictions of renal secretory clearance and pathway-mediated drug interactions during inflammation.

**Visual Abstract:** 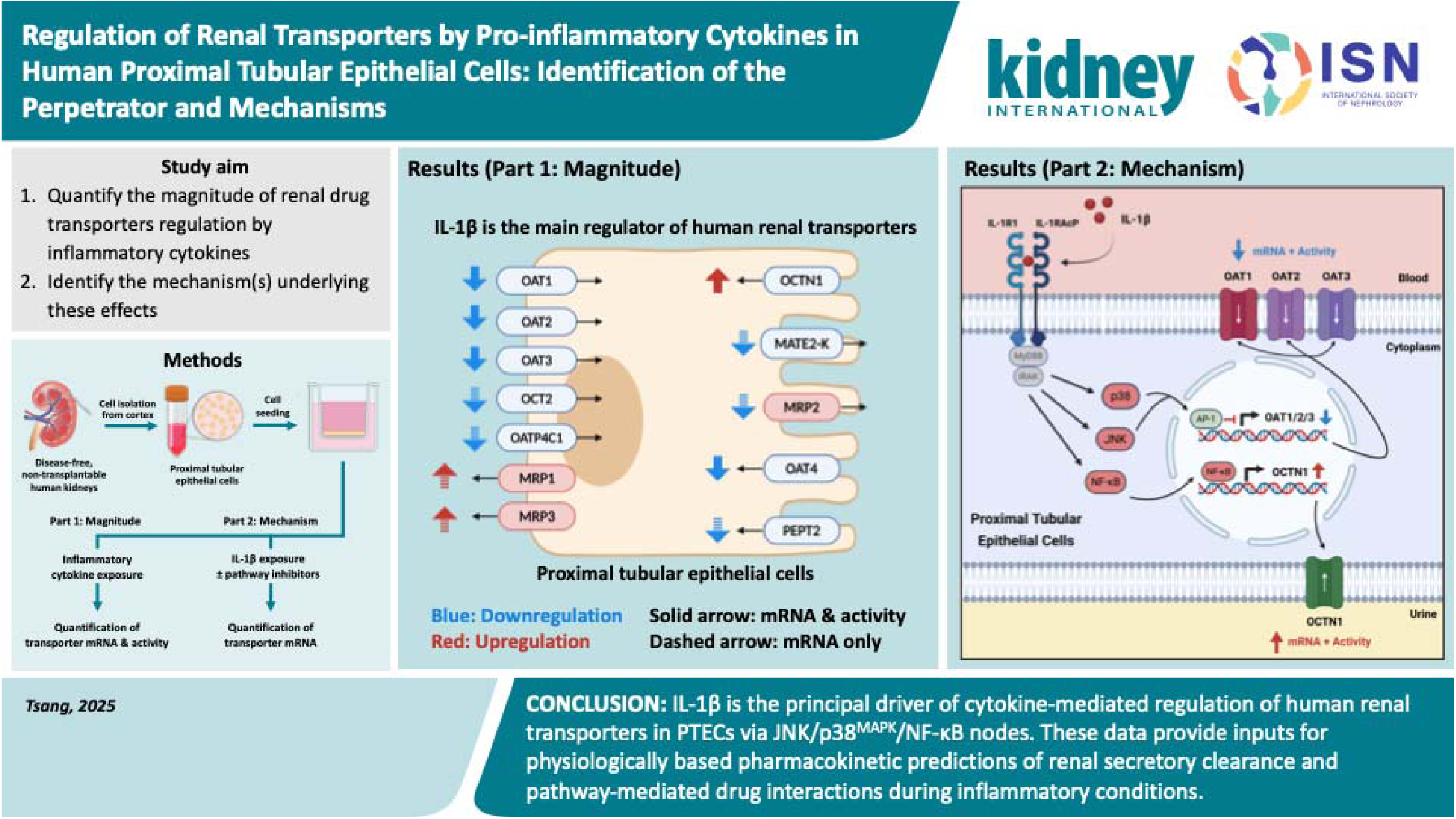

**Translational Statement:** Systemic inflammation increases cytokine concentrations and alters drug pharmacokinetics. Yet, cytokine regulation of renal drug transporters remains poorly defined, even though the kidney clears many anti-infective drugs via active secretion. Using an optimized primary human proximal tubular epithelial cell model that preserves expression and function of major renal transporters, we found that IL-1β is the predominant cytokine that downregulates the mRNA and activity of OAT1–3, OCT2, and OAT4, while upregulating the mRNA and activity of OCTN1. We further showed that IL-1β-driven downregulation of OAT1/3 occurs through JNK signaling, OAT2 through p38^MAPK^, and OCTN1 through NF-κB. These data provide quantitative inputs for physiologically based pharmacokinetic models to predict how inflammation alters renal transporter-mediated drug clearance, informing dose adjustment and risk assessment for disease-drug and drug-drug interactions in patients with inflammatory kidney disease or systemic infections. They also highlight signaling nodes where anti-inflammatory therapies might inadvertently modify renal drug transport.

## Introduction

Conditions such as acute infection, sepsis, and autoimmune disease frequently produce systemic inflammation (1–5). In these settings, circulating cytokines, such as interleukin-6 (IL-6), interleukin-1β (IL□1β), tumor necrosis factor-α (TNF-α), and interferon-γ (IFN-γ), are elevated (**Table 1**). These cytokines regulate drug-metabolizing enzymes and transporters (DMETs), resulting in clinically meaningful changes in drug pharmacokinetics (PK) (1,6–11). While the effects of inflammatory cytokines on hepatic DMETs are well described, their effects on renal transporters are poorly characterized. Yet, many anti□infectives and antivirals rely on renal secretion for clearance (12). Proximal tubular epithelial cells (PTECs) express basolateral uptake transporters, including organic anion transporters 1, 2, and 3 (OAT1–3), organic cation transporter 2 (OCT2), and organic anion transporting polypeptide 4C1 (OATP4C1) (12–14). PTECs also express apical efflux transporters, including multidrug and toxin extrusion proteins 1 and 2-K (MATE1/2-K), multidrug resistance□associated proteins 2 and 4 (MRP2/4), and P□glycoprotein (P-gp) (12–14). Together, these transporters coordinate vectorial drug secretion. Transcriptional or functional perturbation of these transporters can change net drug secretion, reabsorption, and intracellular concentrations in PTECs (12,15).

**Table 1.**
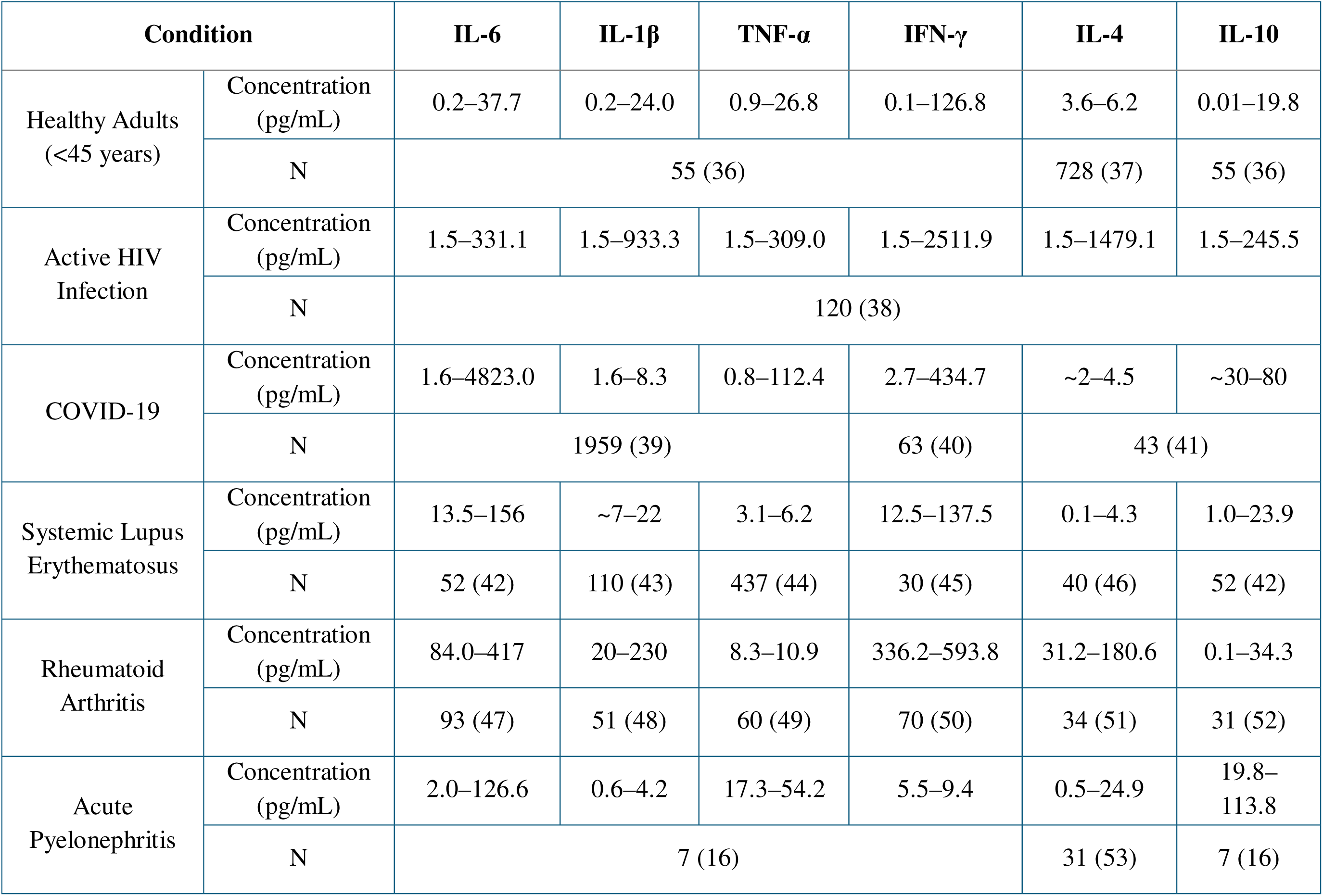
Plasma/serum cytokine concentrations (pg/mL) across inflammatory conditions. Plasma or serum cytokine concentration ranges are shown in representative acute states (e.g., COVID-19, acute pyelonephritis) and chronic states (e.g., HIV infection, systemic lupus erythematosus, rheumatoid arthritis). References are provided in parentheses after the number of subjects (N) for each condition.

| Condition | | IL-6 | IL-1β | TNF-α | IFN-γ | IL-4 | IL-10 |
| --- | --- | --- | --- | --- | --- | --- | --- |
| Healthy Adults<br>(<45 years) | Concentration<br>(pg/mL) | 0.2–37.7 | 0.2–24.0 | 0.9–26.8 | 0.1–126.8 | 3.6–6.2 | 0.01–19.8 |
|  | N | 55 (36) |  |  |  | 728 (37) | 55 (36) |
| Active HIV<br>Infection | Concentration<br>(pg/mL) | 1.5–331.1 | 1.5–933.3 | 1.5–309.0 | 1.5–2511.9 | 1.5–1479.1 | 1.5–245.5 |
|  | N | 120 (38) |  |  |  |  |  |
| COVID-19 | Concentration<br>(pg/mL) | 1.6–4823.0 | 1.6–8.3 | 0.8–112.4 | 2.7–434.7 | ~2–4.5 | ~30–80 |
|  | N | 1959 (39) |  |  | 63 (40) | 43 (41) |  |
| Systemic Lupus<br>Erythematosus | Concentration<br>(pg/mL) | 13.5–156 | ~7–22 | 3.1–6.2 | 12.5–137.5 | 0.1–4.3 | 1.0–23.9 |
|  | N | 52 (42) | 110 (43) | 437 (44) | 30 (45) | 40 (46) | 52 (42) |
| Rheumatoid<br>Arthritis | Concentration<br>(pg/mL) | 84.0–417 | 20–230 | 8.3–10.9 | 336.2–593.8 | 31.2–180.6 | 0.1–34.3 |
|  | N | 93 (47) | 51 (48) | 60 (49) | 70 (50) | 34 (51) | 31 (52) |
| Acute<br>Pyelonephritis | Concentration<br>(pg/mL) | 2.0–126.6 | 0.6–4.2 | 17.3–54.2 | 5.5–9.4 | 0.5–24.9 | 19.8–<br>113.8 |
|  | N | 7 (16) |  |  |  | 31 (53) | 7 (16) |

Only one human study has assessed the impact of inflammation on renal transporter-mediated PK. In pregnant patients with acute pyelonephritis, the secretory clearance of the OAT1/3 probe furosemide was 43% lower during active infection than after resolution of pyelonephritis in a paired design (16). This effect during infection coincided with elevated plasma cytokine concentrations. These observations support inflammation- and cytokine-linked downregulation of renal transporters as a plausible driver of reduced renal secretory clearance of furosemide. Accordingly, these data prompted us to evaluate, in human PTECs, whether cytokines drive this effect, which cytokine is responsible, and the mechanisms by which it does so.

Previously, we tested whether pro-inflammatory cytokines alter renal transporter mRNA in freshly isolated human PTECs cultured on flat plates. Cytokines downregulated OCT2 and OATP4C1 mRNA and upregulated organic cation/carnitine transporter 1 (OCTN1) mRNA (17). However, the instability of OAT1–3 expression in this system precluded reliable quantification of OAT-mediated transport activity. To address this limitation, we cultured primary human PTECs on extracellular matrix (ECM)-coated Transwells and optimized isolation and culture conditions.

In Part 1 of this study, we evaluated the effects of mammalian cell-derived pro-inflammatory (IL-6, IL-1β, TNF-α, IFN-γ) and anti-inflammatory (IL-4, IL-10) cytokines on renal transporter mRNA and on the activity of major renal uptake transporters. Cytokines were tested at 0.1 or 1 ng/mL, individually or in combination (herein referred to as the <u>cytokine</u> <u>cocktail</u>, with each cytokine at 0.1 or 1 ng/mL). These concentrations fall within reported *in vivo* pathophysiological ranges (**Table 1**). IL-4 and IL-10 were included because their effects on renal drug transporters are largely uncharacterized. These cytokines are also relevant to chronic autoimmune diseases and to the resolution phase of acute infections (18,19). We also examined renal drug-metabolizing enzymes (DMEs) and endocytic receptors, which are sparsely covered in the literature. These quantitative readouts are intended to support physiologically based pharmacokinetic (PBPK) modeling and simulations to predict inflammation-dependent changes in the disposition of renally secreted drugs (20–22).

In Part 2, we investigated mechanisms underlying cytokine effects on renal DMETs and endocytic receptors using small-molecule inhibitors. Several cytokine-activated signaling pathways provide plausible mechanisms for transporter regulation in the kidney. IL-1β signals by first binding interleukin-1 receptor type 1 (IL-1R1) and interleukin-1 receptor accessory protein (IL-1RAcP) on the cell surface. Ligand binding recruits myeloid differentiation primary response protein 88 (MyD88) and interleukin-1 receptor-associated kinase 1 and 4 (IRAK1/4), forming the oligomeric myddosome complex. This complex then activates TNF receptor–associated factor 6 (TRAF6) and transforming growth factor-β–activated kinase 1 (TAK1) (23–26). These proteins, in turn, activate mitogen-activated protein kinase (MAPK) cascades, including extracellular signal-regulated kinase (ERK), p38^MAPK^, and c-Jun N-terminal kinase (JNK), as well as the nuclear factor-κB (NF-κB) pathway via the IκB kinase complex (23–26). In contrast, IL-6 signals via glycoprotein (gp130)-associated Janus kinases 1 and 2 (JAK1/2) and tyrosine kinase 2 (TYK2). These kinases phosphorylate signal transducer and activator of transcription 1 (STAT1) and STAT3. The phosphorylated STATs, in turn, dimerize and drive downstream transcription (27–30). Classic signaling requires membrane-bound IL-6 receptor α (mIL-6Rα). Trans-signaling occurs when IL-6 binds soluble IL-6Rα (sIL-6Rα), enabling gp130-positive cells that lack mIL-6Rα to respond (27–30). Pathways activated by both IL-6 and IL-1β intersect with transcriptional programs and nuclear receptors that regulate hepatic DMETs (31–33). Although these mechanisms are better established in hepatic models (1,31,34,35), pathway dependencies of cytokine effects on human renal transporters remain unresolved. Therefore, we tested whether the cytokine-driven changes in transporter mRNA and activity identified in Part 1 are mediated through MAPK and NF-κB signaling pathways. We also tested whether IL-6 classic or trans-signaling produces similar effects in primary human PTECs.

## Materials and Methods

### Chemicals and reagents

A list of chemicals, reagents, and their suppliers is provided in **Supplementary Table S1**.

### Isolation of primary human PTECs

Disease-free, non-transplantable adult human kidneys (glomerular filtration rate > 60 mL/min) were obtained from Organ Procurement Organizations via Novabiosis, Inc. (Durham, NC). Kidneys were deemed unsuitable for clinical transplantation for allocation or quality reasons unrelated to intrinsic kidney disease (e.g., not allocated within the clinical time window, prolonged cold ischemia, or donor serologies incompatible with available recipients). Donor demographics are provided in **Supplemental Table S2**. Kidneys were preserved in sterile University of Wisconsin solution at 4 °C. Cold ischemia time (from vascular clamping to initiation of cell isolation) was less than 36 hours. Under sterile conditions, cortical tissue from kidneys was minced into ∼1 mm^3^ pieces in Hank’s balanced salt solution with Ca² /Mg² (HBSS^+/+^) and digested at 37 °C for 45 min in PES-filtered digestion buffer containing 240 U/mL collagenase IV, 0.6 U/mL Dispase, 25 mM NaHCO_3_, 25 mM HEPES, 3 mM CaCl_2_, and 0.2% BSA in HBSS^+/+^. After digestion, cell suspensions were centrifuged at 300 g for 5 minutes at 4 °C. Pellets were then washed twice with cold Hank’s balanced salt solution without Ca²□/Mg²□(HBSS^−/−^) containing 10 mM EDTA and filtered through 70 µm cell strainers. Cell suspensions were then centrifuged (300 × g, 5 min, 4 °C) and subjected to sequential isotonic Percoll gradients in HBSS^+/+^ (pH 7.4) at 1.02 and 1.07 g/mL at 4 °C (1000 × g for 10 min each step). The resulting 1.07 g/mL supernatants (where PTECs are) were then diluted to <1.02 g/mL with cold HBSS^+/+^ and centrifuged (300 × g, 5 min, 4 °C) to pellet the PTECs. Cell pellets were resuspended in prewarmed PTEC medium (DMEM/F-12 containing 1 g/L D-Glucose, 15 mM HEPES, 25 mM NaHCO_3_, 100 ng/mL EGF, 10 pM triiodothyronine, 100 ng/mL hydrocortisone, 1.72 µM insulin, 68.8 nM transferrin, 38.7 nM sodium selenite, 100 U/mL penicillin, 100 µg/mL streptomycin, and 25 µg/mL of amphotericin B) and seeded on 0.4 µm Transwell inserts coated with 50 µg/cm^2^ of Matrigel (growth factor-reduced, phenol red-free) at a density of 2.5–3.5 × 10^5^ cells/cm^2^. 1 µM A83-01 and 10 µM Y-27632 were included only during the first 24 h after seeding to aid cell differentiation, survival, and attachment. PTECs were cultured for 3 days in a humidified incubator (37 °C, 5% CO_2_), with daily medium changes, before cytokine treatments.

### Cytokine treatments and experimental overview

#### Part 1: Magnitude and directionality of cytokine effects

An experimental overview is shown in **Figure 1**. To quantify the magnitude of cytokine-mediated transporter regulation, PTECs were exposed for 48 h to pro-inflammatory cytokines (IL-6, IL-1β, TNF-α, IFN-γ) and anti-inflammatory cytokines (IL-4, IL-10), individually or in combination (herein referred to as the <u>cytokine cocktail</u>, with each cytokine at 0.1 or 1 ng/mL). Treatments were applied simultaneously to both chambers of the Transwell to mimic systemic exposure (basal) and luminal exposure to cytokine after glomerular filtration (apical). Medium was changed once at 24 h. Following 48 h of incubation, PTECs were harvested for RNA isolation, or used in uptake assays. Basal (OAT1/2/3, OCT2) and apical uptake transporter activity (OAT4, OCTN1) was quantified using selective probe substrates and substrate-inhibitor pairs validated previously (**Supplemental Table S3**). In parallel, mRNA expression of renal DMETs and endocytic receptors was quantified by reverse transcription quantitative PCR (RT-qPCR) with TaqMan probes (**Supplemental Table S4**).

**Figure 1.**
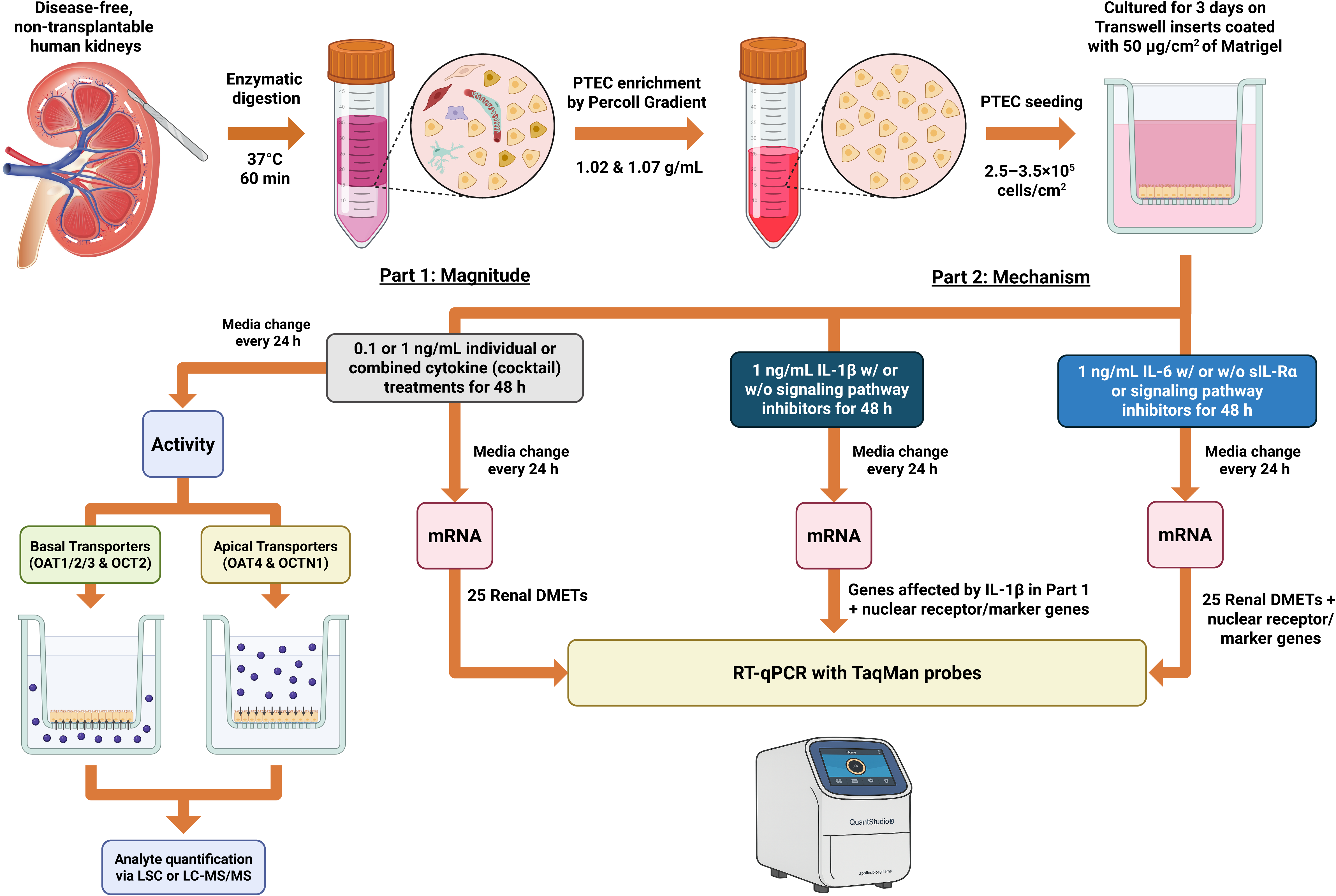
Experimental workflow to assess cytokine-mediated regulation of renal drug transporters, DMEs, and endocytic receptors in primary human PTECs. Disease-free, non-transplantable human kidneys were enzymatically digested (37 °C, 60 min), and PTECs enriched by discontinuous Percoll density gradient (1.02 & 1.07 g/mL). PTECs were then seeded on Matrigel-coated Transwell inserts (2.5–3.5 × 10^5^ cells/cm^2^) and pre-cultured for 3 days. **Part 1:** PTECs were exposed every 24 h, for 48 h, to individual cytokines (IL-6, IL-1β, TNF-α, IFN-γ, IL-4, IL-10) at 0.1 or 1 ng/mL or a <u>cytokine cocktail</u> (each cytokine at 0.1 or 1 ng/mL) in both Transwell chambers. RT-qPCR was performed with TaqMan probes to quantify the mRNA expression of renal DMETs and endocytic receptors. Activity of uptake transporters (OAT1–4, OCT2, OCTN1) was quantified using selective substrates or substrate-inhibitor pairs (**Supplemental Table S3**). **Part 2:** PTECs were exposed to IL-1β (1 ng/mL) in both Transwell chambers every 24 h, for 48 h, with or without small-molecule inhibitors of ERK (10 µM PD98059), p38^MAPK^ (10 µM SB203580), JNK (10 µM SP600125), and IκBα (30 µM PDTC) to test whether MAPK or NF-κB underlie the IL-1β-driven changes observed in **Part 1**. mRNA expression of genes affected by IL-1β in **Part 1** and selected nuclear receptor/inflammatory markers were quantified by RT-qPCR. We also evaluated IL-6 with sIL-6Ra to probe classic versus trans-signaling (57–59) and asked whether trans-signaling can override the lack of effects by IL-6 alone. sIL-6Ra was added only basally to mimic its limited *in vivo* renal filtration due to size (∼50 to 70 kDa). See Methods for additional details. Figure created with BioRender. Kidney illustration was provided by Kayenat Aryeh.

#### Part 2: Mechanistic studies

Because IL-1β produced the most consistent and pronounced effects across transporters in part 1, it was selected for mechanistic evaluation. PTECs were treated with IL-1β (1 ng/mL) for 48 h in the presence or absence of signaling pathway inhibitors in both the apical and basal Transwell chambers. Inhibitors were used either individually or in combination (herein referred to as the <u>inhibitor cocktail</u>), targeting canonical IL-1β signaling cascades: ERK (PD98059, 10 µM), p38^MAPK^ (SB203580, 10 µM), JNK (SP600125, 10 µM), and NF-κB (PDTC, 30 µM) (54,55). The inhibitor cocktail contained all four compounds at the same concentrations used in individual inhibitor conditions. Fresh medium with cytokine (± inhibitors) was replaced after 24 h to maintain activity. At the end of the 48 h treatment, total RNA was extracted, and the mRNA expression of renal DMETs, endocytic receptors, nuclear receptors, and inflammatory markers were quantified by RT-qPCR (**Supplemental Table S4**).

Given IL-6’s role in hepatic transporter regulation (6,56), we also evaluated IL-6 in combination with its soluble receptor sIL-6Rα to distinguish classic from trans-signaling (57–59). PTECs on Transwells were treated for 48 h (media replaced every 24 h) with the following conditions: (i) IL-6 (1 ng/mL) for classic signaling; (ii) IL-6 (1 ng/mL) + sIL□6Rα (100□ng/mL) + inhibitor cocktail for trans-signaling with exogenous IL-6; (iii) sIL□6Rα (100□ng/mL) alone for trans-signaling with endogenous IL-6; and (iv) sIL□6Rα (100□ng/mL) + the inhibitor cocktail to control for sIL-6Rα background and MAPK/NF-κB crosstalk. IL-6 and the inhibitor cocktail were added to both Transwell chambers. sIL-6Rα was only added basally because its large size (∼50–70 kDa) limits renal filtration *in vivo* under disease-free conditions (60). sIL-6Rα concentration (100 ng/mL) was chosen to match its observed *in vivo* plasma concentrations during inflammation (57,61,62). Fresh medium for each condition was replaced after 24 h to maintain activity. After treatments, PTECs were harvested for RNA isolation and RT-qPCR with TaqMan probes. Genes examined are listed in **Supplemental Table S4**.

### Quantification of mRNA expression

At the end of the treatments, total RNA was extracted from PTECs using the PureLink RNA Mini Kit. Samples were treated with DNase I on-column to eliminate genomic DNA and minimize interference with downstream RT-qPCR. RNA concentrations were normalized to the lowest of the batch (within each donor), and cDNA was synthesized using High-Capacity cDNA Reverse Transcription Kit. RT-qPCR was performed with TaqMan probes on a QuantStudio 3 real-time PCR system (Thermo Fisher Scientific, Waltham, MA). A list of TaqMan probes used is provided in **Supplemental Table S4**. Each qPCR reaction contained 10 μL of 2× TaqMan Fast Advanced Master Mix, 1 μL of 20× TaqMan probes, and 9 μL of cDNA diluted in RNase-free water. The thermocycling conditions were 20 s at 95 °C, followed by 40 cycles of 1 s at 95 °C and 20 s at 60 °C.

### Quantification of uptake transporter activity using selective transporter substrates or substrate-inhibitor pairs

Substrates, inhibitors, and their working concentrations are listed in **Supplemental Table S3**. After cytokine treatments, PTECs were washed twice with warm HBSS^+/+^ (basal pH 7.4; apical pH 6.5) and preincubated with inhibitors or vehicle (DMSO) for 15 min at 37 °C. Preincubation buffer was then replaced with warm HBSS^+/+^ containing the relevant substrate (± inhibitor, as indicated) in the basal chamber (pH 7.4, for OAT1, OAT2, OAT3, and OCT2) or the apical chamber (pH 6.5, for OAT4 and OCTN1). Uptake proceeded for 15 min, after which the buffer was aspirated and cells were washed three times with ice-cold HBSS^+/+^. Methods for downstream analyte quantification has been described previously (63). Briefly, for radiolabeled substrates (used for OAT1, OAT2, OCT2, and OCTN1), cells were lysed overnight at room temperature in 1 M NaOH and neutralized with 1 M HCl. Lysates were mixed with Ecoscint liquid scintillation cocktail, and radioactivity was measured on a Tri-Carb B3110TR liquid scintillation counter (PerkinElmer, Waltham, MA). For non-labeled substrates (used for OAT3 and OAT4), cells were lysed in ice-cold 100% acetonitrile containing fexofenadine as the internal standard (50 nM for levocetirizine/OAT4 assays; 250 nM for GCDCA-S/OAT3 assays), diluted 1:1 with water, and quantified by LC–MS/MS as previously described (63). Proteins precipitated in each insert were solubilized in 1 M NaOH overnight at room temperature and neutralized with 1 M HCl the next day. For radioactive assays, total protein was quantified from an aliquot of the same neutralized NaOH lysate used for scintillation counting. For non-radioactive assays, total protein was quantified from the neutralized NaOH-resolubilized protein fraction generated from the acetonitrile precipitate. Protein was measured with the Pierce BCA Protein Assay Kit on a Spark multimode microplate reader (Tecan, Männedorf, Switzerland). Absolute substrate uptake was normalized to the BCA-measured protein mass in the aliquot used for analyte quantification (pmol/mg).

### Cytokine and sIL-6Rα quantification by ELISA

Cytokine concentrations in apical and basal media collected after the first 24 h of treatment were measured using Human ELISA kits from Proteintech (IL-6, IL-1β, TNF-α, IFN-γ, IL-4, IL-10) and R&D Systems (sIL-6Rα), following manufacturers’ instructions. Cytokine concentrations in medium at the start of treatment were also sampled (“initial,” immediately after cytokine addition). Samples were diluted in the kit-specific assay diluent to fall within the assays’ linear range (IL-6: 15.6–1000 pg/mL; IL-1β: 3.9–250 pg/mL; TNF-α: 31.25–2000 pg/mL; IFN-γ: 15.6–1000 pg/mL; IL-4: 15.6–1000 pg/mL; IL-10: 7.8–500 pg/mL; sIL-6Rα: 31.2–2,000 pg/mL). Absorbance was read on a Spark multimode microplate reader (Tecan, Männedorf, Switzerland).

### Data and Statistical Analysis

Relative mRNA expression versus vehicle (2^−ΔΔCt^) was calculated using the 2^−ΔΔCt^ method, where ΔC_t_ (cycle threshold) = C_t_ of the gene of interest − C_t_ of GAPDH (housekeeping gene), and ΔΔC_t_ = ΔC_t_ of treated sample − average ΔC_t_ of the vehicle-treated samples (64). GAPDH remained stable within donors across conditions at a fixed cDNA input for RT-qPCR (**Supplemental Figure S1**). C_t_ values above 37 were excluded due to poor signal-to-noise. For mechanism studies, because pathway inhibitors altered baseline expression, values were normalized by taking the ratio of 2^−ΔΔCt^ in cytokine + inhibitor samples to the respective inhibitor-only (cytokine-free) condition.

Active transporter-mediated uptake (pmol/mg) was computed within each condition as absolute substrate uptake amount in non-inhibited wells minus uptake in inhibited wells. Relative active uptake to vehicle control was then expressed as the ratio of active transporter uptake in treated group to that in vehicle-treated group.

For ELISA, standard curve absorbances (kit-provided standards) were plotted against nominal concentrations and fitted with the four-parameter logistic model. Sample concentrations were then back-calculated in GraphPad Prism 10.2.1 (GraphPad Software, La Jolla, CA) and multiplied by the corresponding dilution factors.

Specific statistical tests are detailed in the Results and the figure legends. In brief, comparisons to vehicle-treated samples (**Figures 2–4, 6, and Supplemental Figures S4–S6**) used two-way analysis of variance (ANOVA) with Dunnett’s multiple comparisons correction (row factor: concentration; column factor: treatment). Mechanism studies (**Figures 5, 7, and S8)** used repeated measures one-way ANOVA with Geisser-Greenhouse correction (for sphericity) and Dunnett’s multiple comparisons correction (or Šídák’s multiple comparisons for pre-selected pairs, see **Figure 8**). All data analyses were performed on GraphPad Prism 10.2.1 (GraphPad Software, La Jolla, CA).

**Figure 2.**
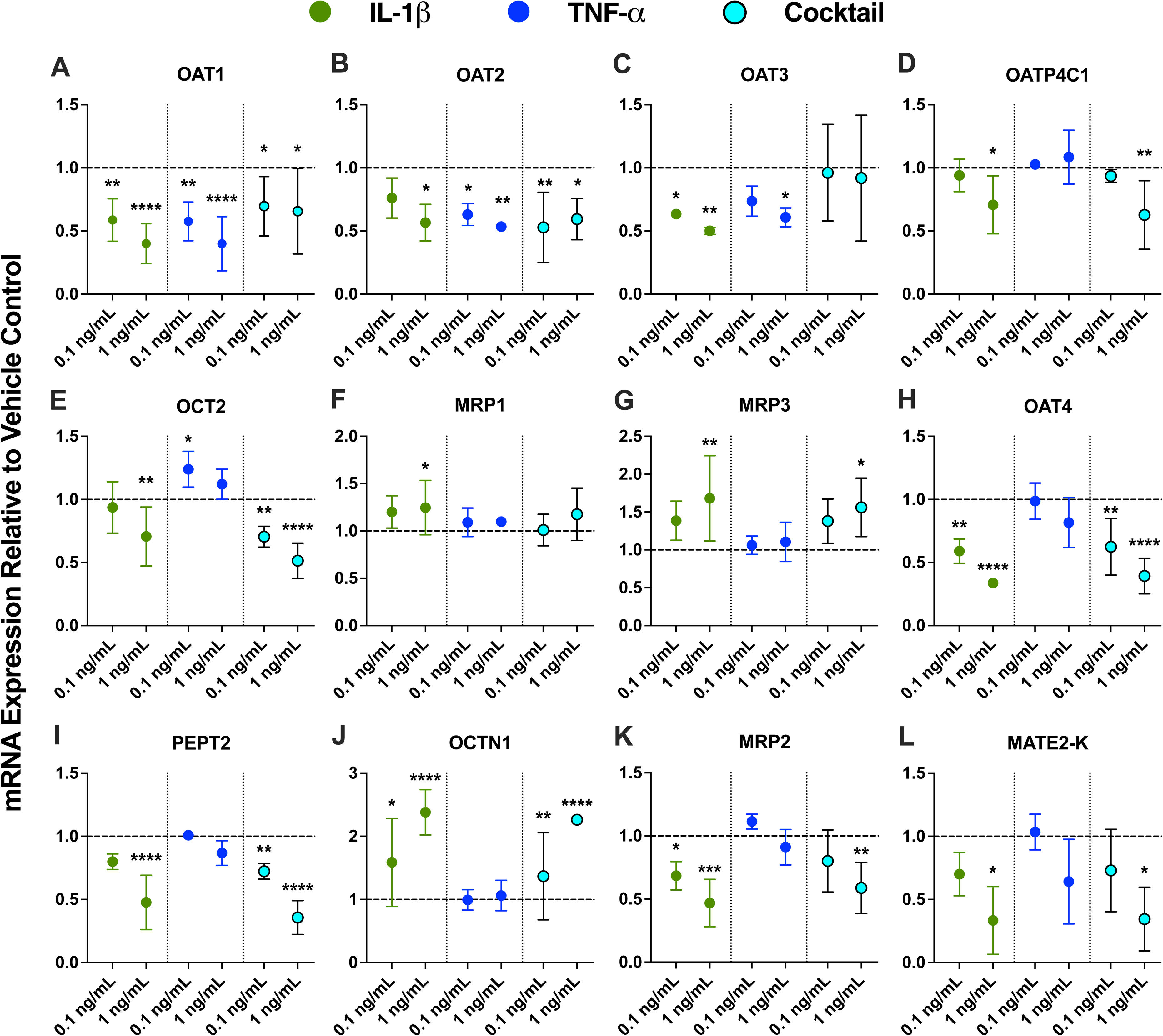
IL-1β was the main modulator of renal transporter mRNA in primary human PTECs, with TNF-α exerting weaker effects. PTECs on Transwells were exposed to individual cytokines (IL-6, IL-1β, TNF-α, IFN-γ, IL-4, and IL-10) or the cytokine cocktail at 0.1 and 1 ng/mL in both chambers every 24 h for 48 h. Only data for IL-1β (green), TNF-α (blue), and the cytokine cocktail (teal) are plotted in this figure because they produced the most pronounced effects. Data for IL-6, IFN-γ, IL-4, and IL-10, which produced smaller or sporadic effects, are shown in **Supplemental Figure S4**. Panels: **basal uptake transporters** ([**A**] OAT1, [**B**] OAT2, [**C**] OAT3, [**D**] OATP4C1, [**E**] OCT2), **basal efflux transporters** ([**F**] MRP1, [**G**] MRP3), **apical uptake transporters** ([**H**] OAT4, [**I**] PEPT2, [**J**] OCTN1), **apical efflux transporters** ([**K**] MRP2, [**L**] MATE2-K). mRNA was normalized to GAPDH and expressed relative to vehicle control (0.1% DPBS; dashed line at y = 1). Across donors, IL-1β produced the largest, concentration-dependent changes: downregulation of OAT1–3, OCT2, OATP4C1, OAT4, PEPT2, MRP2, and MATE2-K mRNA and upregulation of MRP1/3 and OCTN1 mRNA. TNF-α downregulated the mRNA of OAT1–3 to a similar extent as IL-1β but had little effect on other transporters. Effects of the cytokine cocktail were generally similar to those of IL-1β (except for OAT3). Data are mean ± SD from three donors, each quantified in technical triplicate. Statistical significance was assessed using two-way ANOVA with Dunnett’s multiple comparisons (*p ≤0.05, **p<0.01, ***p<0.001, ****p <0.0001). Additional results for other DMETs and endocytic receptors are provided in **Supplemental Figure S4**.

**Figure 3.**
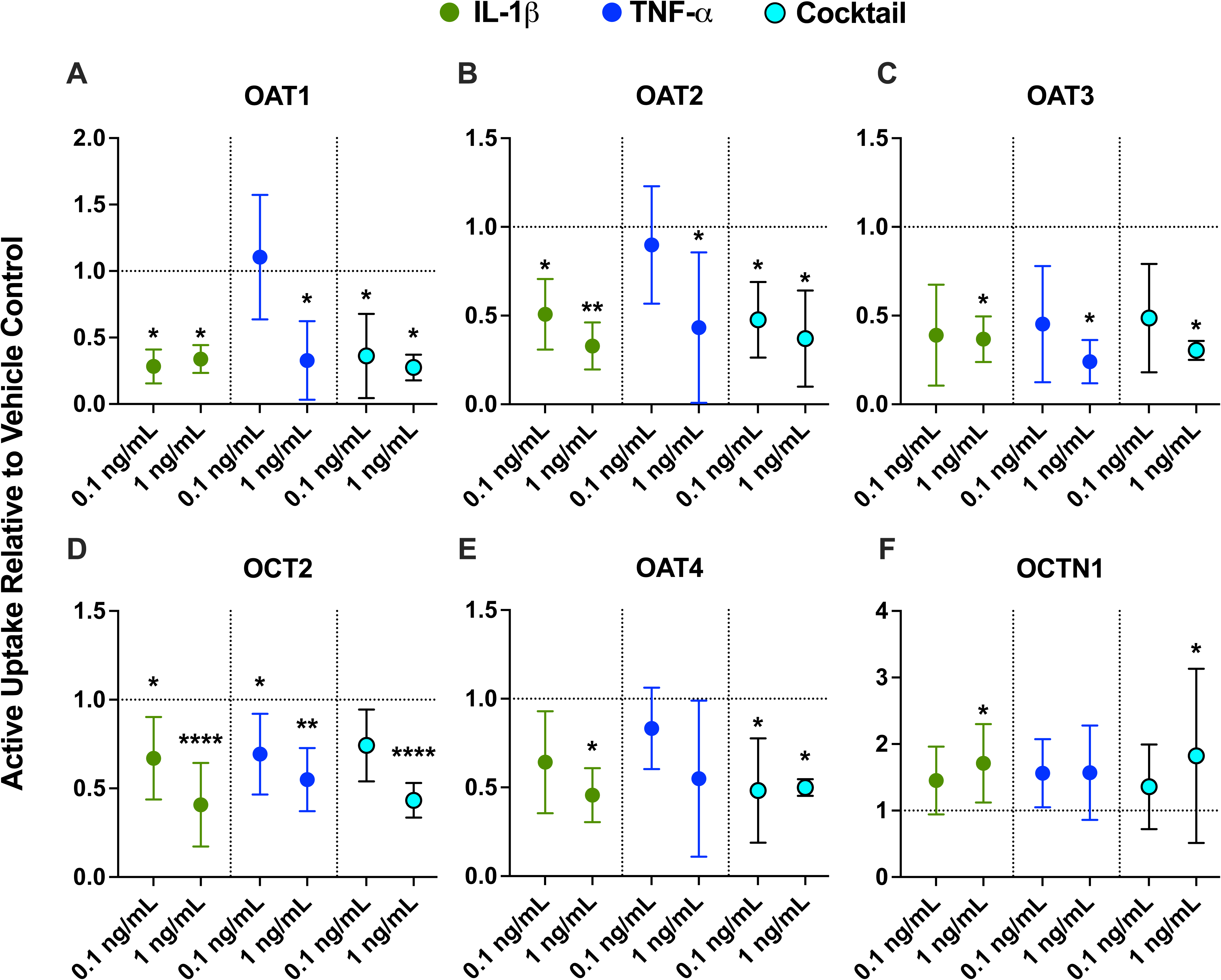
Renal uptake transporter activity in PTECs was modulated mainly by IL-1β and TNF-α, with overall patterns largely mirroring the mRNA responses. PTECs on Transwells were exposed to individual cytokines (IL-6, IL-1β, TNF-α, IFN-γ, IL-4, and IL-10) or the cytokine cocktail at 0.1 and 1 ng/mL in both chambers for 48 h (media replaced every 24 h). Only data for IL-1β (green), TNF-α (blue), and the cytokine cocktail (teal) are plotted in this figure because they produced the most pronounced effects. Data for IL-6, IFN-γ, IL-4, and IL-10, which produced smaller or sporadic effects, are shown in **Supplemental Figure S5**. Activity of uptake transporters ([**A**] OAT1, [**B**] OAT2, [**C**] OAT3, [**D**] OCT2, [**E**] OAT4, [**F**] OCTN1) is presented as the fraction of active uptake (determined by normalizing transporter-selective substrate uptake in the absence of inhibitors to that measured in the presence of inhibitors [**Supplemental Table S3**]) relative to vehicle-treated controls (0.1% DPBS; horizontal dashed line at y = 1). Similar to the mRNA results, IL-1β (and partially TNF-α) produced the broadest and largest changes in renal uptake transporter activity. Effects of the cytokine cocktail generally mirrored those of IL-1β. Data are mean ± SD from three donors (each quantified in triplicate). Statistical significance (*p≤0.05, **p<0.01, ****p<0.0001) was assessed using two-way ANOVA with Dunnett’s multiple comparisons.

**Figure 4.**
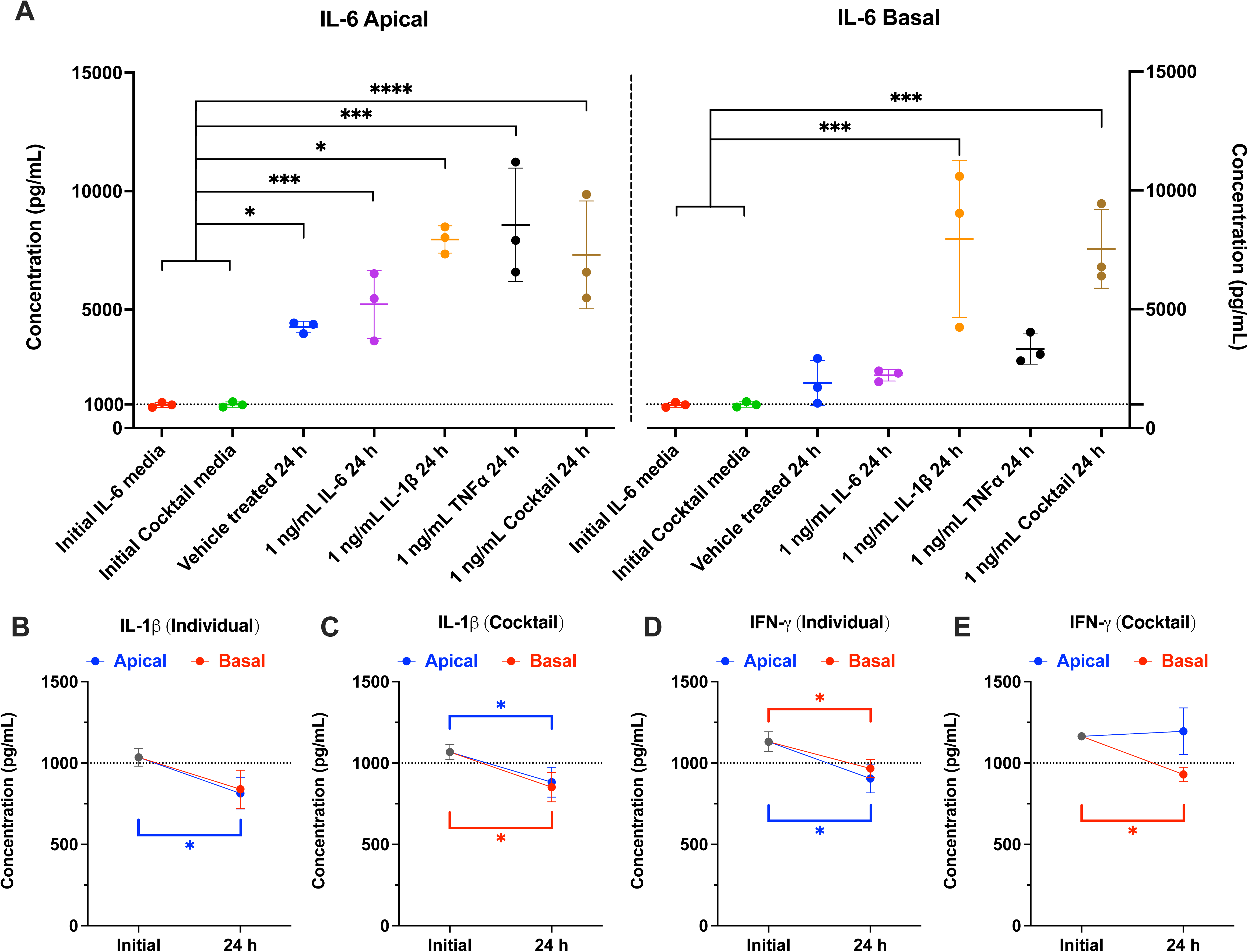
Endogenous IL-6 production by PTECs and cytokine concentration in media over 24 h of exposure. Cytokines (IL-6, IL-1β, TNF-α, IFN-γ, IL-4, and IL-10) were quantified in culture media by ELISA at the start of treatment (“initial”, cytokine-containing treatment media before cell contact) and after the first 24 h of 1 ng/mL exposure (media was changed every 24 h; 48 h of total treatment time). Apical and basal compartments were sampled. Data are ± SD from three donors (denoted by points, each quantified in triplicate). Dotted lines denote the initial nominal 1 ng/mL concentration. Statistical significance (*p≤0.05, ***p<0.001, ****p<0.0001) was assessed using one-way ANOVA with Dunnett’s multiple comparisons. (**A**) IL-6 accumulated in vehicle-treated cultures, above the initial 1 ng/mL spike, and increased further with 1 ng/mL of IL-1β or TNF-α treatment, consistent with endogenous and IL-1β/TNF-α stimulated secretion of IL-6 by PTECs. Exogenous cytokine concentrations were largely stable over 24 h (>80%), with small but significant decreases for IL-1β (**B, C**) and IFN-γ (**D, E**); TNF-α, IL-4, and IL-10 concentrations showed no significant change (**Supplemental Figure S6**).

**Figure 5.**
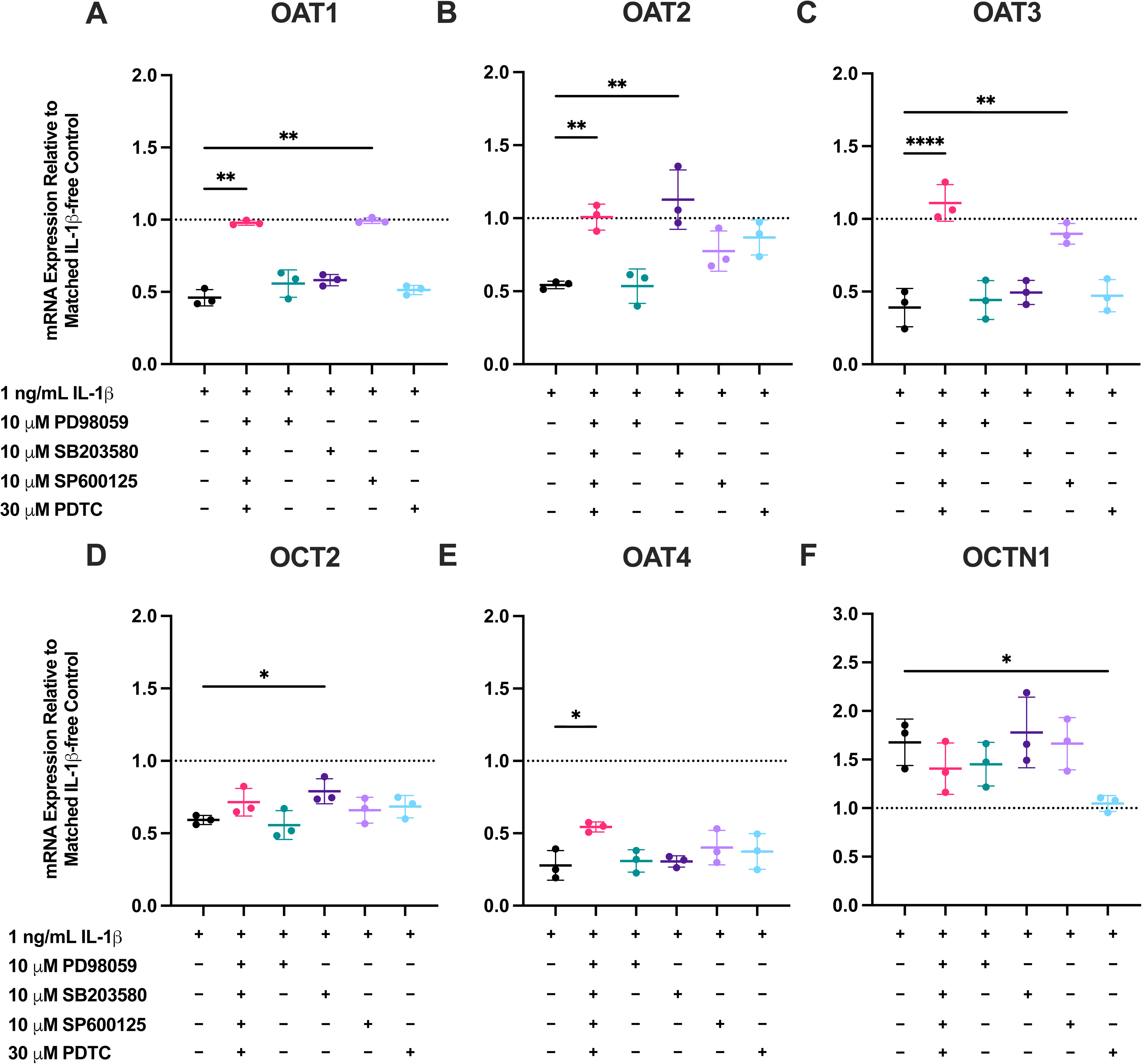
MAPK and NF-κB blockade mitigate IL-1β-driven transcriptional changes of key renal uptake transporters in PTECs. Primary human PTECs were treated for 48 h (media replaced every 24 h) with 1 ng/mL IL-1β in the absence or presence of pathway inhibitors (i.e., PD98059 [ERK, 10 µM], SB203580 [p38^MAPK^, 10 µM], SP600125 [JNK, 10 µM], and PDTC [NF-κB, 30 µM]), either individually or as a cocktail, in both chambers of Transwells. Panels: (**A**) OAT1, (**B**) OAT2, (**C**) OAT3, (**D**) OCT2, (**E**) OAT4, and (**F**) OCTN1. Expression is shown relative to the respective IL□1β□free controls within each inhibitor condition (dotted line = 1), as these inhibitors alone without cytokines affected the mRNA expression of transporters (**Supplemental Figure S7**). Thus, IL-1β + inhibitor cocktail data are normalized to inhibitor cocktail-only controls, and IL-1β + individual inhibitor data are normalized to the corresponding inhibitor-only controls. Data are mean ± SD from three donors (each quantified in triplicate). Statistical significance was assessed using repeated-measures one-way ANOVA with Dunnett’s multiple comparisons against the corresponding IL-1β-free baselines (*p≤0.05, **p<0.01, ****p <0.0001). The inhibitor cocktail completely reversed IL-1β-mediated repression of OAT1–3. JNK blockade alone (SP600125) fully restored OAT1/3, whereas for OAT2, p38^MAPK^ inhibition (SB203580) significantly blunted the effect of IL-1β. OCTN1 induction by IL-1β was significantly reduced by NF-κB inhibition. OCT2 effects were modest overall, with only p38^MAPK^ inhibition reaching significance. OAT4 was rescued by the cocktail, whereas no single inhibitor reproduced the cocktail’s effect. Data for additional DMETs and endocytic receptors tested are shown in **Supplemental Figures S7** and **S9**. Quantification of marker gene mRNA expression assessing pathway engagement by inhibitors is shown in **Supplemental Figure S8.**

## Results

### PTEC isolation, identity, and experimental overview

Primary human PTECs were isolated from six disease-free, adult, non-transplantable donor kidneys (**Supplemental Table S2**) and seeded on Matrigel-coated Transwell inserts (**Figure 1**). GAPDH was stable across treatment conditions within each donor and was used for mRNA normalization (**Supplemental Figure S1**). Cultures retained a proximal tubule phenotype, with higher relative mRNA expression of aquaporin 1 (AQP1; proximal tubule marker) compared to AQP2 (cortical collecting duct marker) and sodium-chloride cotransporter (NCC; distal tubule marker) (**Supplemental Figure S2**). Baseline transporter mRNA expression in untreated cultures after 5 days is provided in **Supplemental Table S5**. Compared with flat-plate culture, Transwells maintained similar OAT1/3 mRNA expression but greater OAT2 mRNA expression (OAT2 was undetectable in flat-plate cultures after 5 days) (**Supplemental Figure S3A**). Activity of OAT1–3 was measurable over the experimental timeline (**Supplemental Figure S3B**).

### IL-1β was the main transcriptional regulator of renal DMETs and endocytic receptors in primary human PTECs

Across the six cytokines, IL-1β produced the broadest and largest concentration-dependent mRNA changes in renal DMETs and endocytic receptors (**Figure 2 and Supplemental Figure S4**). TNF-α reproduced the OAT1–3 suppression by IL-1β with similar magnitude, but had little effect on other genes (**Figure 2**). Effects of the cytokine cocktail generally mirrored those of IL-1β, except for OAT3 and UDP-glucuronosyltransferase 1A9 (UGT1A9) (**Figure 2 and Supplemental Figure S4**). IFN-γ caused only modest changes in renal DMET mRNA. IL-6, IL-4, and IL-10 had little to no significant effects on the mRNA of renal DMETs and endocytic receptors (**Supplemental Figure S4**). For clarity, only data for IL-1β, TNF-α, and the cytokine cocktail are shown in **Figure 2**, while data for all the cytokines are shown in **Supplemental Figure S4**.

At 0.1 ng/mL, IL 1β significantly downregulated the mRNA of OAT1 (by 41.3%), OAT3 (by 36.5%), OAT4 (by 40.9%), MRP2 (by 31.5%), cytochrome P450 2B6 (CYP2B6) (by 28.1%), and UGT2B7 (by 33.3%), while upregulating the mRNA of OCTN1 (by 59.1%) and CYP3A5 (by 60.7%) (**Figure 2 and Supplemental Figure S4**). At 1 ng/mL, IL 1β significantly downregulated the mRNA expression of OAT1 (by 60.0%), OAT2 (by 43.3%), OAT3 (by 49.9%), OATP4C1 (by 29.3%), OCT2 (by 29.4%), OAT4 (by 66.2%), peptide transporter 2 (PEPT2) (by 52.4%), MRP2 (by 66.6%), MATE2-K (by 53.2%), CYP2B6 (by 48.8%), UGT2B7 (by 37.7%), cubilin (CUBN) (by 30%), and megalin (LRP2) (by 46.1%) (**Figure 2 and Supplemental Figure S4**). IL 1β at 1 ng/mL also significantly upregulated the mRNA expression of MRP1 (by 24.6%), MRP3 (by 66.2%), OCTN1 (by 138%), and CYP3A5 (by 122%) (Figure 2 and Supplemental Figure S4).

TNF-α reproduced IL-1β effects but only for OAT1–3 (**Figure 2A–C**). At 0.1 ng/mL, TNF-α significantly downregulated the mRNA of OAT1 (by 42.4%) and OAT2 (by 36.9%) (**Figure 2A–B**). TNF-α at 0.1 ng/mL also significantly but modestly upregulated OCT2 mRNA (by 23.9%), but this effect was not seen at the higher concentration (1 ng/mL) (**Figure 2E**). At 1 ng/mL, TNF-α downregulated the mRNA expression of OAT1 (by 60.1%), OAT2 (by 46.6%), and OAT3 (by 39.2%) (**Figure 2A–C**).

IFN-γ, at 1 ng/mL, significantly but modestly downregulated OATP4C1 (by 29.3%), PEPT2 (by 38.1%), sodium-glucose cotransporter (SGLT2) (by 25.2%), and UGT1A9 (by 51.6%) (**Supplemental Figure S4**). The cytokine cocktail also produced qualitative effects that mirror those by IL 1β, except for OAT3 (no significant effects) and UGT1A9 (downregulation mirrors IFN-γ) (**Figure 2 and Supplemental Figure S4**). IL-6 had no significant effects on renal DMET and endocytic receptor mRNA, except for a modest upregulation of OCT2 mRNA (by 24%) at 0.1 ng/mL (**Supplemental Figure S4**). IL-4 and IL-10 had no significant effects, except for upregulation of MRP2 (by 32%) and P-gp (by 39.7%) by IL-4 at 1 ng/mL (**Supplemental Figure S4**). mRNA expression of other transporters tested (i.e., OCTN2, urate transporter 1 (URAT1), breast cancer resistance protein (BCRP), MATE1) were not significantly affected by the inflammatory cytokines tested (**Supplemental Figure S4**).

### IL-1β- and TNF-α-driven changes in renal uptake transporter activity mirrored transcriptional changes

Similarly to the mRNA results, IL-1β produced the broadest and largest changes in renal uptake transporter activity. IL-1β exposure significantly reduced the activity of basal OAT1–3 and OCT2 and apical OAT4, while inducing the activity of OCTN1 (**Figure 3**). TNF-α similarly reduced activity of basal OAT1–3 and OCT2 but did not significantly alter OAT4 and OCTN1 activity (**Figure 3**). Effects of the cytokine cocktail generally mirrored those of IL-1β (**Figure 3**). IFN-γ, IL-4, and IL-10 caused limited, transporter activity changes (**Supplemental Figure S5**). IL-6 had no significant effect on the activity of renal uptake transporters (**Supplemental Figure S5**).

Compared to the mRNA data (**Figure 2**), activity data had considerably greater inter-donor variability, which could be due to the comparatively lower sensitivity of functional assays versus RT-qPCR. Across all three donors, IL-1β at 0.1 ng/mL significantly reduced the activity of OAT1 by 71.7%, OAT2 by 49.2%, and OCT2 by 33.0% (**Figure 3A–B, D**). At 1 ng/mL, IL-1β significantly reduced the activity of OAT1 by 66.2%, OAT2 by 67.1%, OAT3 by 63.3%, and OCT2 by 59.2% (**Figure 3A–D**). At 0.1 ng/mL, TNF-α significantly reduced OCT2 activity by 30.7% (**Figure 3D**). At 1 ng/mL, TNF-α significantly reduced the activity of OAT1 by 67.2%, OAT2 by 57.1%, OAT3 by 76%, and OCT2 by 45.1% (**Figure 3A–D**). These effects were largely concentration-dependent, except for OAT1 where a slight greater reduction in activity was observed for IL-1β at 0.1 ng/mL compared to 1 ng/mL **(Figure 3A)**. For apical transporters, 1 ng/mL of IL-1β significantly reduced the activity of OAT4 by 54.3% and induced the activity of OCTN1 by 71.1% (**Figure 3E and F**). The effects of the cytokine cocktail were largely the same as IL-1β. IFN-γ also significantly downregulated OAT1 activity by 62.5% (**Supplemental Figure S5A**). IL-4 at 0.1 ng/mL led to a modest but significant reduction of OCT2 activity (by 23.5%) (**Supplemental Figure S5D**). IL-10 modestly but significantly reduced OCT2 activity (by 22.7%) at 0.1 ng/mL (**Supplemental Figure S5D)**. At 1 ng/mL, IL-10 significantly induced OAT1 activity by 81.3% and reduced OAT2 activity by 55.2% (**Supplemental Figure S5A and B)**. While the magnitude of OAT1 activity upregulation by IL-10 was greater than that of reduction by IL-1β and TNF-α, the cytokine cocktail results more closely reflected the reduction (**Supplemental Figure S5A**). IL-6, similar to the mRNA data, had no effects on uptake transporter activity (**Supplemental Figure S5**).

Overall, the directionality of cytokine-driven changes in the activity of renal uptake transporters agrees with the mRNA data (**Figure 2**), though they differ in magnitude. For instance, TNF-α downregulated the mRNA expression of OAT3 by 39.2% (**Figure 2C**), while it reduced OAT3 activity by 76% (**Figure 3C**).

### Cytokine stability and endogenous IL-6 production by PTECs

To verify PTEC exposure to the cytokines and determine whether PTECs release cytokines, we quantified cytokine concentrations in apical and basal media by ELISA at the start of treatment (initial, before cell contact) and after the first 24 h of exposure. Because cytokine stability at 0.1 and 1 ng/mL was comparable in pilot experiments, we proceeded with the measurement of cytokine stability at 1 ng/mL. Even in vehicle-treated cultures, IL-6 in apical culture supernatant after 24 h significantly increased to concentrations above 1 ng/mL, indicating endogenous secretion by PTECs (**Figure 4A**). IL 1β and cytokine cocktail treatment further significantly increased IL 6 in both compartments (apical > basal) to similar concentrations (**Figure 4A**). TNF-α treatment also significantly increased IL-6 concentrations in the apical compartment (**Figure 4A**). These results are consistent with prior observations in PTECs in which IL-6 is endogenously produced and IL-1β/TNF-α stimulate IL-6 secretion by PTECs (65).

Concentrations of other exogenously added cytokines are largely stable over 24 h and remained near the measured initial treatment concentrations (>80%), with small but significant decreases for IL 1β (individual and cocktail conditions; **Figure**□**4B–C**) and IFN γ (**Figure**□**4D–E**). TNF α, IL 4, and IL 10 concentrations showed no significant changes over 24 h (**Supplemental Figure S6**). The modest losses observed here for cytokines, except for IL-6, indicate that the effective exposure approximated the intended 1 ng/mL (or 0.1 ng/mL). Therefore, the observed mRNA and activity changes (**Figures 2 and 3**) can be attributed to ∼1 ng/mL (or 0.1 ng/mL) cytokine treatments rather than cytokine instability-driven reduced concentrations.

### MAPK/NF-κB blockade reverses IL-1β-mediated transporter mRNA changes with node-specific dependencies

Because IL-1β was the dominant suppressor of basal OAT1–3 (mRNA and activity) as well as many other genes tested in Part 1, we next probed its signaling mechanism. PTECs from three additional donors (**Supplemental Table S2**) were co-treated, every 24 h, for 48 h with 1 ng/mL IL-1β in the absence or presence of pathway inhibitors targeting ERK (10 µM PD98059), p38^MAPK^ (10 µM SB203580), JNK (10 µM SP600125), or NF-κB (30 µM PDTC), individually or as a four-inhibitor cocktail, in both the apical and basal Transwell chambers. Because the inhibitors themselves altered transporter mRNA expression (**Supplemental Figure S7**), data were normalized and compared to the respective inhibitor-only, IL□1β□free control, within each condition (dotted line□=□1), allowing for a true rescue-of-effect interpretation. Quantification of marker genes assessing pathway engagement by inhibitors is shown in **Supplemental Figure S8**.

The effects of IL-1β on renal DMET and endocytic receptor mRNA in PTECs from Donors 4–6 recapitulated those observed in Donors 1–3 (**Figure 2 and Supplemental Figures S4 and S7**). Across Donors 4–6, the inhibitor cocktail significantly and completely reversed the IL-1β-mediated repression of the mRNA expression of OAT1, OAT2, and OAT3, returning expression to normalized baseline (**Figure 5A–C**). Parsing single nodes revealed JNK inhibition alone significantly restored OAT1 and OAT3 (**Figure 5A and C**), whereas p38^MAPK^ inhibition significantly blunted OAT2 suppression by IL-1β (**Figure 5B**). For OAT2, the effects of JNK or NF-κB blockade were not significant (**Figure 5B**). These patterns indicate that IL□1β suppresses the basolateral OATs via distinct MAPK pathways (JNK for OAT1/3 and p38^MAPK^ for OAT2), with convergence and partial redundancy revealed by the full cocktail rescue. While the magnitudes of rescue varied across donors, the directionality of rescue was consistent.

For OCT2, the effect of the inhibitors was overall modest, as only p38^MAPK^ inhibition reached significance (**Figure**□**5D**), suggesting the involvement of other IL 1β signaling pathways. For OAT4, simultaneous MAPK/NF-κB inhibition by the inhibitor cocktail significantly but modestly restored expression, whereas no single inhibitor reproduced the effect (**Figure 5E**). For OCTN1, IL□1β-driven mRNA induction was significantly attenuated only by NF-κB inhibition.

Rescue of other genes is shown in **Supplemental Figure S7** (vehicle normalized) and **Supplemental Figure S9** (respective inhibitor group normalized). OATP4C1 downregulation was significantly rescued by the inhibitor cocktail, SB203580 (p38^MAPK^), and SP600125 (JNK) (**Supplemental Figures S7 and S9A**). IL-1β effects on MATE2-K, MRP2, PEPT2, CYP3A5, and CYP2B6 were significantly reversed by only the inhibitor cocktail (**Supplemental Figures S7 and S9D–H**). MRP1/3 induction was attenuated by all inhibitor conditions but only in a significant manner for PD98059 (ERK), SP600125 (JNK), and PDTC (NF-κB) to below their respective baseline (i.e., downregulation) (**Supplemental Figures S7 and S9B–C**). This suggests that IL-1β potentially regulates MRP1/3 via parallel nodes that produce opposite effects (up vs downregulation). UGT2B7 downregulation was significantly attenuated by the inhibitor cocktail, as well as SP600125 and PDTC (**Supplemental Figures S7 and S9I**).

### Pathway blockade lowers basal and IL□1β–stimulated IL□6 secretion

To functionally validate pathway inhibitor activity, we measured PTEC-secreted IL-6 in apical and basal media after 24 h of a 48 h treatment period. Cells were treated with 1 ng/mL IL-1β with or without pathway inhibitors used singly or as a cocktail in both Transwell chambers. Endogenous IL-6 secretion was detectable in vehicle-treated cells, and IL-1β significantly induced IL-6 secretion (**Figure 6**). The inhibitor cocktail significantly reduced both basal and IL-1β-stimulated IL-6. Single inhibitors partially attenuated IL-1β-stimulated IL-6 secretion, and had no apparent effects on endogenous IL-6 concentrations in vehicle-treated cells. These functional data corroborate with IL-6 mRNA expression in PTECs under each inhibitor condition (**Supplemental Figure S8**). Notably, induction of IL-6 mRNA was only partially reversed by the inhibitor cocktail despite complete suppression of IL-6 protein secretion (**Figure 6**). This highlights the importance to functionally validate the magnitude of IL-6 stimulation. Across conditions, IL-6 concentrations appear higher in apical chamber (**Figure 6A**) than that in basal (**Figure 6B**).

**Figure 6.**
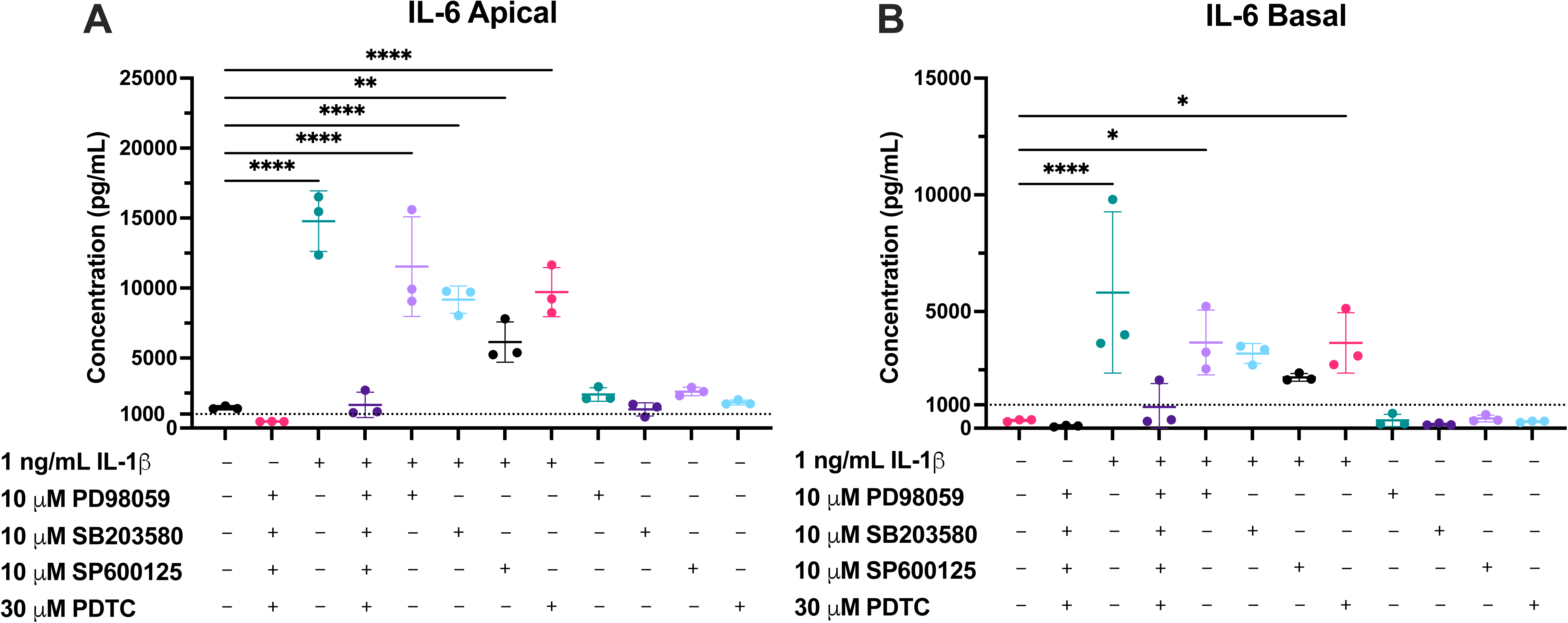
Pathway blockade modulates endogenous and IL□1β–stimulated IL□6 secretion from PTECs. Primary human PTECs cultured on Transwells were treated for 48 h (media replaced every 24 h) with 1 ng/mL IL-1β in the absence or presence of pathway inhibitors (i.e., PD98059 [ERK, 10 µM], SB203580 [p38^MAPK^, 10 µM], SP600125 [JNK, 10 µM], and PDTC [NF-κB, 30 µM]), either individually or as a cocktail, in both chambers of Transwells. At the 24 h timepoint, (**A**) apical and (**B**) basal media were collected and IL-6 concentrations quantified by ELISA. Data are mean (in pg/mL) ±□SD of three donors (denoted by points, each donor quantified in technical triplicate). Statistical significance was assessed using one-way ANOVA with Dunnett’s multiple comparisons against the vehicle control (*p≤0.05, **p<0.01, ****p <0.0001). Endogenous IL-6 secretion was detectable in vehicle-treated group. IL-1β robustly induced IL-6 secretion, and this induction was strongly suppressed by the inhibitor cocktail. Each single inhibitor partially attenuated IL-1β-stimulated secretion of IL-6, but not to the extent of the cocktail. In the absence of IL-1β, single inhibitors had little to no effect on endogenous secretion. Overall, across conditions, IL-6 concentrations are higher in the apical chamber than in the basal chamber.

With the inhibitor cocktail, IL-1β-stimulated IL-6 in media fell markedly. Yet GAPDH-normalized transporter mRNA (e.g., OAT1/2/3) remained higher than that in vehicle-treated PTECs (**Supplemental Figure S7**). Thus, under these conditions, the IL-1β effect is unlikely to be mediated, or confounded, by the IL-6/JAK/STAT cascade. Instead, it is consistent with direct dependence of the MAPK/NF κB axis as shown in **Figure 5**.

### IL□6 classic and trans□signaling do not recapitulate IL□1β effects on transporter mRNA

IL-6 trans-signaling was further probed with the addition of sIL 6Rα. Endogenous IL-6 secretion and the reduction of IL-6 secretion by the inhibitor cocktail were confirmed via ELISA (**Figure 8A**). The concentrations of sIL□6Rα across conditions significantly decreased by ∼60–70% over 24 h, suggesting cellular internalization of the IL-6-sIL□6Rα complex via gp130 (**Figure 8B**) (66–68). Adding sIL□6Rα did not significantly lower IL-6 concentrations over 24 h after accounting for inhibitor effects (**Figures 4, 6 and 8A**). This aligns with the binding kinetics of IL-6/sIL□6Rα showing that most circulating IL-6 remains free at physiologic IL-6/sIL 6Rα ratios (61,62).

Across donors, IL-6 alone (exogenous + endogenous) and IL-6□+□sIL-6Rα + inhibitors showed no significant differences in transporter mRNA after normalization to the respective controls (**Figure 7A–F**). sIL-6Rα alone, with or without inhibitors, also had no significant effects, indicating that IL-6 signaling did not alter renal transporter mRNA expression via either JAK/STAT or MAPK/NF-κB. In contrast, IL-6 (1 ng/mL) + sIL□6Rα (100□ng/mL) + inhibitors significantly downregulated PEPT2 and UGT2B7. However, this effect was absent when PTECs were exposed to 100□ng/mL sIL□6Rα alone, suggesting differences in exogenous and endogenous IL-6 signaling activity for these two genes. No significant effects were observed for other transporters, DMEs, and endocytic receptors (**Supplemental Figures S7 and S10**).

**Figure 7.**
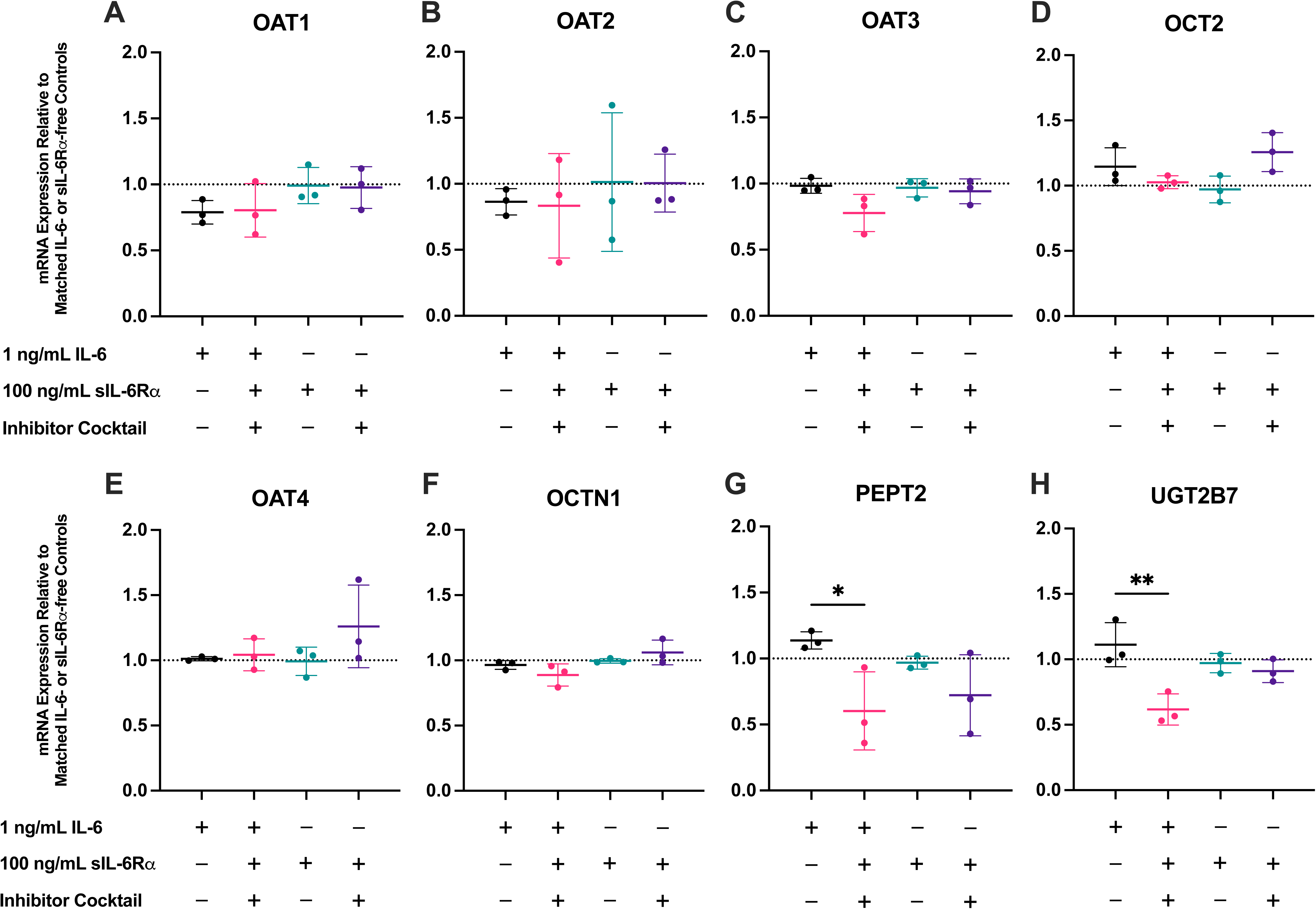
Exogenous IL-6, alone or with sIL-6Rα (trans-signaling), does not significantly alter renal transporter mRNA expression in PTECs. Primary human PTECs cultured on Transwells were treated for 48 h (media replaced every 24 h) with 1 ng/mL IL-6, 1 ng/mL IL-6 + 100□ng/mL sIL□6Rα + the inhibitor cocktail (consist of PD98059 [ERK, 10 µM], SB203580 [p38^MAPK^, 10 µM], SP600125 [JNK, 10 µM], and PDTC [NF-κB, 30 µM]), 100□ng/mL sIL□6Rα alone (to probe the trans-signaling with endogenously secreted IL-6), and 100□ng/mL sIL□6Rα + the inhibitor cocktail (for background sIL-6Rα effects and inhibition of MAPK/NF-κB crosstalk). IL-6 and the inhibitor cocktail were added to both apical and basal chamber of Transwells, while sIL-6Rα was only added to the basal chamber of Transwells as apical sIL-6Rα exposure is not physiologically relevant owing to its greater molecular weight (∼50–70 kDa) and minimal renal filtration *in vivo*. Panels: (**A**) OAT1, (**B**) OAT2, (**C**) OAT3, (**D**) OCT2, (**E**) OAT4, (**F**) OCTN1, (**G**) PEPT2, and (**H**) UGT2B7. Expression is shown relative to the respective IL□6 free or sIL□6Rα-free controls within each inhibitor condition (dotted line□=□1), as these inhibitors alone without cytokines affected the mRNA expression of transporters (**Supplemental Figure S7**). Data are mean ±□SD from three donors (denoted by points, each quantified in triplicate). Statistical significance was assessed using repeated-measures two-way ANOVA with Dunnett’s multiple comparisons against the corresponding IL-6-free or□sIL-6Rα-free baselines (*p≤0.05, **p<0.01, ****p <0.0001). Across transporters, exogenous IL-6 (classic signaling) and IL-6□+□sIL-6Rα (trans-signaling) produced no significant transcriptional changes. Statistically significant downregulation of PEPT2 and UGT2B7 was observed when 1 ng/mL exogenous IL-6, 100□ng/mL sIL□6Rα, and the inhibitor cocktail were added in combination. However, this was not observed when PTECs were exposed to 100□ng/mL sIL□6Rα alone, suggesting differences in signaling activity between exogenous and endogenous IL-6. Data for additional transporters, DMEs, and endocytic receptors tested are in **Supplemental Figures S7** and **S10**. Pathway engagement/marker genes assessing inhibitor activity are shown in **Supplemental Figure S9**.

**Figure 8.**
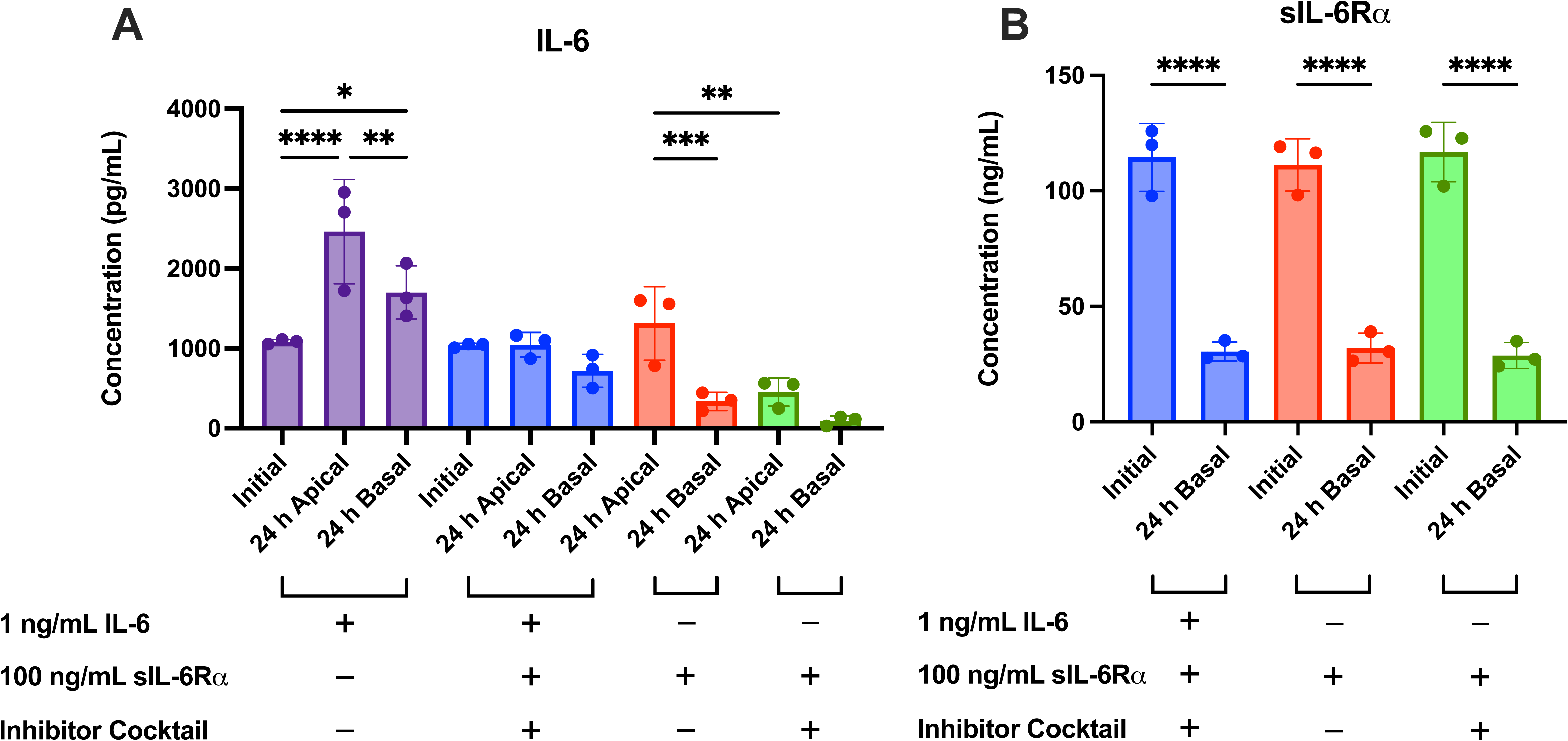
IL-6 and sIL-6Rα concentrations over 24 h with signaling pathway blockade. Primary human PTECs cultured on Transwells were treated for 48 h (media replaced every 24 h) with 1 ng/mL IL-6, 1 ng/mL IL-6 + 100□ng/mL□sIL 6Rα + the inhibitor cocktail, 100□ng/mL sIL□6Rα, or 100□ng/mL sIL□6Rα + the inhibitor cocktail. IL-6 and the inhibitor cocktail were added to both Transwells chambers, while sIL-6Rα was only added basally. Apical and basal media were sampled at the time of addition (“Initial”) and after 24 h, and the concentrations of (A) IL-6 and (B) sIL-6Rα were measured by ELISA. In groups without exogenous IL-6, the inhibitor cocktail reduced endogenous IL-6 to near background. Addition of sIL-6Rα did not further lower IL-6 concentration when the effect of the inhibitor cocktail was accounted for, indicating substantial free IL-6 at steady state. sIL-6Rα significantly declined over 24 h across conditions, consistent with complex formation and receptor-mediated consumption/turnover. The inhibitor cocktail did not meaningfully alter the concentrations of sIL-6Rα over 24 hours, despite lowering the production of endogenous IL-6. Bars show mean ± SD of three donors (denoted by points, each quantified in technical triplicate). Statistical significance (*p≤0.05, **p<0.01, ****p <0.0001) was assessed using repeated-measures one-way ANOVA with Šídák’s multiple comparisons correction between pre-selected pairs (i.e., 0 h vs. 24 h, apical vs basal).

## Discussion

The key advances of this work are summarized below. First, we established a human primary PTEC Transwell system that preserves proximal tubule phenotype long enough to quantify DMET/endocytic receptor mRNA and uptake transporter activity (OAT1–4, OCT2, OCTN1). Second, we quantified cytokine effects on renal DMET/endocytic receptor mRNA and uptake transporter activity in a primary human model using selective substrates/inhibitors. We identified IL-1β as the principal modulator of renal DMET/endocytic receptor mRNA and uptake transporter activity in primary human PTECs. Third, we verified cytokine exposure over the treatment period with ELISA, revealing minimal catabolism of exogenous cytokines and confirming endogenous IL-6 secretion by PTECs. Fourth, we assigned mechanism to the IL-1β-driven suppression of OAT1–3, identifying JNK as a mediator for OAT1/3 and p38^MAPK^ for OAT2 (**Figure 9**). Fifth, we distinguished the effects of IL-6 classic versus trans-signaling while accounting for endogenous IL-6 secretion by PTECs. Collectively, our findings define the magnitude and mechanism of renal transporter regulation by inflammatory cytokines and can be used to inform PBPK models to predict renal secretory clearance during inflammation.

**Figure 9.**
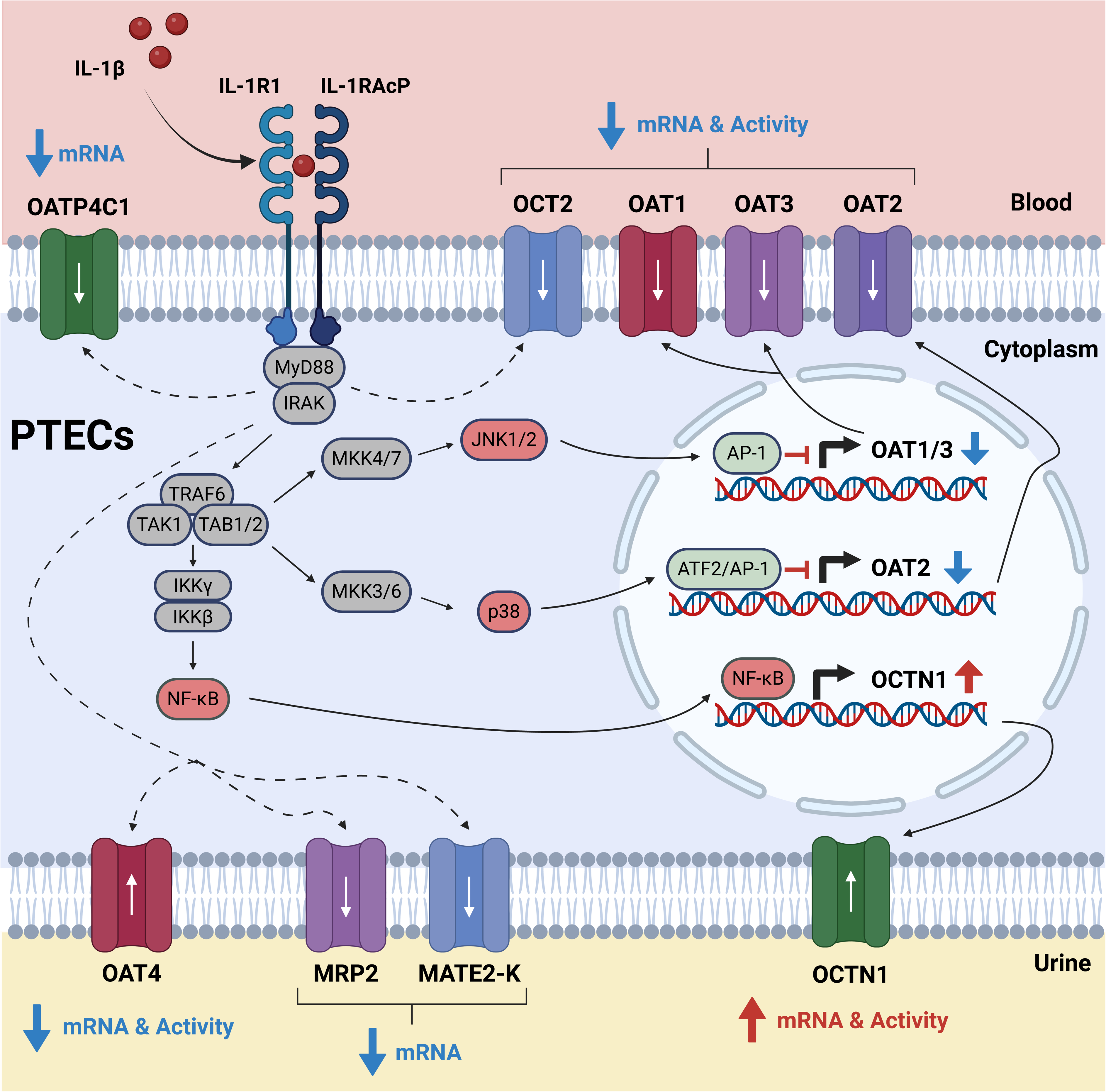
Proposed mechanism by which IL-1β regulates renal transporters in human PTECs. **IL**-1β binds IL-1R1/IL-1RAcP and assembles a myddosome of MyD88, IRAK4, and IRAK1. IRAK4 phosphorylates IRAK1, and together with TRAF6 builds K63-linked ubiquitin scaffolds that activate TAK1-TAB1/2. TAK1 then phosphorylates (i) IKKβ, which phosphorylates IκBα and targets it for degradation, freeing NF-κB to translocate to the nucleus; (ii) MKK4/7, which then dual-phosphorylate JNK1/2; and (iii) MKK3/6, which then dual-phosphorylate p38. Active JNK phosphorylates c-Jun, forming nuclear AP-1 complexes. p38 phosphorylates ATF2 and downstream chromatin kinases, which complexes with c-Jun to form AP-1. These transcription factors (AP-1, ATF2/AP-1, NF-κB) subsequently regulate renal transporter expression (JNK-dependent OAT1/3 downregulation; p38-dependent OAT2 downregulation; NF-κB-dependent OCTN1 upregulation). IL-1β also downregulates OCT2 (mRNA and activity), OAT4 (mRNA and activity), MRP2 (mRNA only), MATE2-K (mRNA only), and OATP4C1 (mRNA only). These effects were observed experimentally but the precise upstream nodes remain to be fully resolved. Blue ↓ = IL-1β-mediated downregulation; Red ↑ = IL-1β-mediated upregulation. Black solid arrows trace canonical signaling pathways. Red-labeled nodes (JNK1/2, p38, NF-κB) highlight key mediators implicated in this study. Downstream transcription factors (AP-1, ATF2/AP-1, NF-κB) indicate the putative regulatory controller. The adjacent blue/red arrows show the net expression change. Dashed arrows denote unresolved pathways that require further investigation. White arrows inside transporters denote physiological substrate transport direction.

Maintaining robust OAT activity in primary human PTECs has been difficult for us and others (17,69). Commercially available primary human PTECs do not express OATs at the mRNA and protein level due to long propagation time across multiple passages (70). While fresh PTEC isolation from donor kidneys partially addresses this issue, OAT1–3 mRNA expression declines rapidly with no measurable activity in conventional flat-plate cultures (17). A primary human PTEC Transwell model that maintains OAT1/3 activity (71) also did not maintain OAT1/3 activity in our hands, when tested with five kidneys. Here, we addressed this limitation by optimizing viability and assay conditions end-to-end. Tissue processing used short and gentle enzymatic digestion. This was followed by Percoll gradient to enrich PTECs, which have a buoyant density of ∼1.036–1.052 g/mL (72). Cells were seeded at near-confluence density onto Matrigel-coated Transwell inserts, which supplies essential ECM proteins (i.e., laminin and collagen IV) and promotes tight junction formation. A brief three-day pre-culture included Y-27632 (Rho-associated protein kinase inhibitor) to reduce dissociation-induced apoptosis and improve early attachment and A83-01 (transforming growth factor-β receptor inhibitor) to blunt TGF-β-driven epithelial-to-mesenchymal transition (73–76). Transport assays preserved physiological compartmentalization (basal pH at 7.4, apical pH at 6.5). Together, this workflow maintained transporter phenotype (mRNA and activity) long enough to quantify cytokine effects across donors.

Across donors, IL-1β was the principal modulator of transporter mRNA and activity (**Figure 2**) in a concentration-dependent manner. At 0.1 ng/mL, IL-1β significantly downregulated the mRNA of OAT1, OAT3, OAT4, and MRP2, while upregulating OCTN1 (**Figure 2**). At the higher tested concentration (1 ng/mL), IL-1β amplified its effects on the above transporters while additionally downregulating OAT2, OATP4C1, OCT2, PEPT2, and MATE2-K, while upregulating basal efflux transporters MRP1/3 (**Figure 2**). At 1 ng/mL, TNF-α had similar effects as IL-1β in terms of both directionality and magnitude, but only for OAT1–3. For the major renal uptake transporters (i.e., OAT1–4, OCT2, OCTN1), these IL-1β-driven mRNA changes, as well as TNF-α driven mRNA changes of OAT1–3, were directionally correlated with activity changes by functional assays using selective substrates or selective substrate-inhibitor combinations (**Figure 3**) (63). However, the mRNA and activity changes differed in magnitude. For instance, TNF-α downregulated the mRNA expression of OAT3 by 39.2% (**Figure 2C**), while it reduced OAT3 activity by 76% (**Figure 3C**). Since protein turnover rates are typically longer than mRNA turnover rates (77), this suggests that cytokines could be post-transcriptionally regulating renal transporters. Importantly, ELISA results showed more than 80% stability in the medium of exogenously added cytokines over 24 hours. Thus, the observed effects reflect near-nominal concentrations rather than cytokine-depletion driven reduced concentrations (**Figures 4 and S6**).

Collectively, our results are consistent with an inflammatory cytokine-driven reprogramming of vectorial transport in human proximal tubule epithelium. IL-1β (with TNF-α concordant for OAT1–3) suppressed basal substrate uptake, limiting epithelial entry of circulating organic anions and cations. Concomitant suppression of apical OAT4 and PEPT2 would be expected to reduce luminal reabsorption of their substrates, while decrease in MRP2 and MATE2-K indicate a global reduction of the apical efflux capacity for their substrates. In contrast, induction of basolateral MRPs suggests a redirection of conjugated metabolites and oxidative-stress products toward the interstitium or blood. Notably, while OCTN1 is an uptake transporter for ergothioneine, it has been reported to mediate epithelial acetylcholine export (78). Our data suggest that IL-1β could activate downstream cholinergic, α7-nicotinic acetylcholine receptor-dependent anti-inflammatory signaling in PTECs (79–81). Overall, these results suggest that inflammation minimizes PTEC intracellular solute accumulation and dampens pro-inflammatory signaling at the expense of reduced renal secretory clearance of endogenous and exogenous substrates of the affected transporters.

The magnitude of IL-1β-driven OAT1/3 downregulation at 0.1 ng/mL (**Figures 2 and 3**) agrees quantitatively, in terms of directionality and magnitude expected for a reduction in basal uptake, with what we observed *in vivo* during active pyelonephritis (16). Notably, measured plasma cytokines in that cohort were toward the low end of pathophysiological ranges (<0.1 ng/mL), and plasma IL-1β concentrations were not significantly elevated during active pyelonephritis. Together, these observations suggest that plasma cytokine concentrations during infection/inflammation may underreport the cytokine exposure experienced by PTECs in the interstitial microenvironment and at the basolateral membrane. This gap emphasizes the need for quantitative systems pharmacology linking circulating cytokines to predicted interstitial and luminal cytokine concentrations in the human kidney. This can be followed by PBPK modeling and simulation using transporter-specific directionality and magnitude derived from human PTECs to predict the effect of inflammation on the PK of renally secreted drugs.

Our mechanism studies placed JNK upstream of IL-1β–driven OAT1/3 repression and p38^MAPK^ upstream of OAT2 (**Figure 5A–C**). OCT2 and OAT4 only showed weak sensitivity to MAPK/NF-κB blockade. Partial reversal of OCT2/OAT4 downregulation with MAPK/NF-κB blockade suggests additional regulatory nodes (e.g., phosphatidylinositol 3-kinase/protein kinase B [PI3K/AKT]) or post-transcriptional mechanisms (**Figure 5D–E**) (25). NF-κB inhibition alone attenuated OCTN1 mRNA induction, consistent with prior reports in MH7A fibroblast-like synoviocytes (82). However, the inhibitor cocktail did not further reduce OCTN1 mRNA induction, suggesting antagonistic crosstalk in which MAPK blockade partially relieves a parallel restraint on OCTN1 induction and offsets NF-κB inhibition (**Figure 5F**). These findings have several clinical implications. First, they enable risk reasoning for disease-drug interactions and drug-drug interactions (DDI) that are not obvious from other well-characterized regulatory pathways (e.g., PXR- and CAR-mediated induction of CYP3A4 and P-gp). Drugs that chronically suppress JNK or p38^MAPK^ signaling in the kidney would be expected to relieve IL-1β-like repression of OAT1–3 less effectively than a broad anti-inflammatory that also dampens upstream cytokine production. Second, because inhibition of these pathways by small molecules alone upregulated renal transporter mRNA expression (**Supplemental Figure S7**), drugs that chronically target these pathways could increase the secretory clearance via OATs. A similar strategy has been explored for JNK inhibitor bosutinib to mitigate radiation toxicity (83). Indeed, many immunomodulators and targeted oncology agents intersect with MAPK or NF-κB signaling (84–88), and systemic inflammatory diseases often co-occur with these therapies (89–92). Importantly, any indirect DDI would be expected only when basolateral OAT1/3 uptake is the rate-determining step for vectorial secretion (93). In that scenario, pathway modulators that reverse IL-1β repression will proportionally increase OAT1/3-mediated drug uptake and net secretory clearance of OAT substrates, even without direct transporter inhibition.

A common counter-hypothesis is that IL-6 signaling in the kidney is the main driver of transporter changes. IL-6 signaling via JAK/STAT has been implicated in a number of fibrotic kidney diseases, such as lupus nephritis, diabetic nephropathy, and chronic kidney disease (94–96). While this was plausible based on our Part 1 findings, data from further testing in Part 2 do not support that view under the conditions tested. PTECs secreted IL-6 endogenously, and IL-1β and TNF-α further increased IL-6 in both Transwell compartments (**Figure 4**), which was thought to confound the lack of effects from exogenous IL-6 (**Figures 2 and 3**). Yet when IL-6 secretion was abolished by the inhibitor cocktail, exogenous IL-6 did not alter the mRNA expression of renal transporters (**Figure 7**). Addition of sIL-6Rα to enable trans-signaling also did not recapitulate IL-1β effects. The inhibitor cocktail lowered endogenous IL-6 secretion while reversing IL-1β-driven transporter changes, indicating that the IL-1β phenotype in PTECs is not mediated by the canonical JAK/STAT cascade of IL-6 in this context. While this does not exclude that IL-6 can engage MAPK or NF-κB in other renal cell types or disease states, it indicates that any IL-6 contribution to transporter regulation in PTECs is likely contingent on the same nodes that IL-1β dominates. These findings align with the data observed for IL-1β and with the lack of effect from IL-6 + sIL-6Rα at concentrations that consumed a substantial fraction of sIL-6Rα over 24 hours (**Figures 7 and 8**).

Several limitations of this study should be acknowledged. First, dissection of mechanisms relied on small-molecule inhibitors. Although we controlled for inhibitor-dependent baselines and verified pathway engagement with ELISA and RT-qPCR, small molecules can have off-target actions and may not cleanly separate branches where crosstalk is substantial. Nonetheless, selectivity of the inhibitors and their respective concentrations were confirmed based on the literature (97–101), and we do not expect significant crosstalk to have occurred during the experiments. Future experiments incorporating genetic perturbations in longer-lived *in vitro* kidney systems (e.g., organoids or tubule-on-a-chip) (13,69,102,103) would further strengthen the causal inferences in this study. Second, we only quantified the effect of cytokines on the activity of the major uptake transporters. Functional testing of other transporters affected by IL-1β at the mRNA level (i.e., OATP4C1, MRP1–3, MATE2-K) would require transporter-selective substrates. Efflux transporters would also require validated transcellular flux protocols to separate passive diffusion and basal uptake from true apical efflux. This was beyond the scope of the present study and will be pursued in future studies. Third, our cytokine exposures were limited to 48 hours at 0.1 and 1 ng/mL. Inflammatory milieus in patients are time-varying, involve additional mediators, and likely produce interstitial cytokine concentrations that diverge from plasma. A complete concentration-time characterization of IL-1β (and TNF-α for OAT1–3) in this Transwell system would provide the dose-response information needed for translational PBPK modeling. Finally, the cytokine cocktail consisted of equal concentrations for each cytokine to enable controlled comparisons. However, *in vivo* cytokine stoichiometry varies by disease stage and tissue compartment. Future studies should vary cytokine composition and concentrations to test for non-additive or synergistic/antagonistic interactions and potential switching of the main perpetrator.

In summary, we established an optimized primary human PTEC isolation and culture method that overcomes the rapid loss of OAT activity typical of primary cultures. Using this system, we showed that IL-1β is the principal driver of cytokine-mediated renal transporter regulation. JNK and p38^MAPK^ mediate basal OAT1/3 and OAT2 repression by IL-1β, respectively, and NF-κB is required for OCTN1 induction. IL-6, despite robust endogenous production, did not recapitulate the effect of IL-1β through classic or trans-signaling under the conditions tested. These findings provide a mechanistic, cytokine exposure-verified dataset for predicting changes in renal secretory clearance during inflammation using translational PBPK modeling and highlight pathway nodes where indirect renal transporter-mediated interactions may arise.

## Acknowledgements

We thank Dr. Lawrence Lash (Wayne State University) for his guidance in primary PTEC culture, Dale Whittington for his assistance with LC–MS analysis, and Kayenat Aryeh for her kidney illustration in Figure 1.

## Data Availability Statement

The authors declare that all the data supporting the findings of this study are available within the paper and its Supplemental Data.

## Supplementary Material

### Supplemental File

Supplemental Figure S1. GAPDH stability across donors and conditions.

Supplemental Figure S2. Relative mRNA expression of renal cortical cell markers.

Supplemental Figure S3. OAT mRNA expression and activity in primary human PTECs cultured on Transwell inserts versus flat-plates.

Supplemental Figure S4. Full heatmap summary of cytokine effects on the mRNA expression of renal drug transporters, drug metabolizing enzymes, and endocytic receptors.

Supplemental Figure S5. Renal uptake transporter activity in PTECs was modulated mainly by IL-1β and TNF-α.

Supplemental Figure S6. Cytokine stability in apical and basal media over the first 24 h of exposure to PTECs on Transwells (ELISA).

Supplemental Figure S7. Full heatmap summary of the regulation of renal DMET and endocytic receptor mRNA expression by IL-1β ± signaling pathway inhibitors and IL-6 trans-signaling with sIL-6Rα (vehicle-normalized).

Supplemental Figure S8. GAPDH-normalized mRNA expression of nuclear receptors and pathway-engagement markers in PTECs after treatments with IL-1β ± pathway inhibitors or with IL-6 classic/trans-signaling related conditions, relative to vehicle control.

Supplemental Figure S9. Effect of pathway inhibitors on IL-1β–mediated regulation of renal DMET mRNA (matched-inhibitor normalized).

Supplemental Figure S10. Full heatmap summary of the effect of IL-6 classic and trans-signaling on renal DMET and endocytic receptor mRNA (matched-inhibitor–normalized).

Supplemental Table S1. Chemicals and Reagents.

Supplemental Table S2. PTEC donor demographics.

Supplemental Table S3. Substrates and inhibitors used for uptake assays in primary human PTECs.

Supplemental Table S4. TaqMan Assay IDs

Supplemental Table S5. Baseline GAPDH-normalized mRNA expression of renal transporters in vehicle-treated PTECs on Transwells after 5 days in culture.

## Author Contributions Statement

Y.P.T. wrote the manuscript; Y.P.T., Q.M., E.J.K, and J.D.U. designed the research; Y.P.T. and K.W. performed the research; Y.P.T. and K.W. analyzed the data; K.W., E.J.K, and J.D.U. reviewed and edited the manuscript.

## Footnotes

-These data were presented in part as posters and podium presentations at the 2025 Drug Metabolism Gordon Research Conference (GRC) and the 14^th^ International Meeting of International Society for the Study of Xenobiotics (ISSX).

## Abbreviations

ATF2: Activating transcription factor 2
ANOVA: Analysis of variance
AP-1: Activator protein 1
AQP1: Aquaporin 1
AQP2: Aquaporin 2
BCA: Bicinchoninic acid
BCRP: Breast cancer resistance protein
BSA: Bovine serum albumin
CAR: Constitutive androstane receptor
cDNA: Complementary DNA
c-Jun: Transcription factor Jun
CUBN: Cubilin
DDI: Drug-drug interaction
DMEM/F-12: Dulbecco’s modified eagle medium/nutrient mixture F-12
DME: Drug metabolizing enzymes
DMET: Drug metabolizing enzymes and transporters
DMSO: Dimethyl sulfoxide
DNase I: Deoxyribonuclease I
DPBS: Dulbecco’s phosphate-buffered saline
DUSP1: Dual specificity phosphatase 1
DUSP6: Dual specificity phosphatase 6
EGF: Epidermal growth factor
ELISA: Enzyme-linked immunosorbent assay
ERK: Extracellular signal–regulated kinase
ECM: Extracellular matrix
GAPDH: Glyceraldehyde-3-phosphate dehydrogenase
GCDCA-S: Glycochenodeoxycholic acid sulfate
gp130: Glycoprotein 130
HBSS+/+: Hank’s balanced salt solution with Ca²□/Mg²□
HBSS−/−: Hank’s balanced salt solution without Ca²□/Mg²□
HNF1α: Hepatocyte nuclear factor 1 alpha
HNF4α: Hepatocyte nuclear factor 4 alpha
IFN-γ: Interferon-gamma
IκBα: NF-κB inhibitor alpha
IKK: NF-κB inhibitor alpha kinase
IL-1β: Interleukin-1 beta
IL-1R1: Interleukin-1 receptor type 1
IL-1RAcP: Interleukin-1 receptor accessory protein
IL-4: Interleukin-4
IL-6: Interleukin-6
IL-10: Interleukin-10
IRAK: Interleukin-1 receptor-associated kinase
JAK/STAT: Janus kinase/signal transducer and activator of transcription
JNK: c-Jun N-terminal kinase
LC–MS/MS: Liquid chromatography–tandem mass spectrometry
LRP2: Megalin or low density lipoprotein-related protein 2
LSC: Liquid scintillation counting
MAPK: Mitogen-activated protein kinase
MATE: Multidrug and toxin extrusion protein
MKK: Mitogen-activated protein kinase kinase
mIL-6Rα: Membrane-bound interleukin-6 receptor alpha
MEK: MAPK/ERK kinase
MRP: Multidrug resistance–associated protein
MyD88: Myeloid differentiation primary response protein 88
NCC: Sodium-chloride cotransporter
NF-κB: Nuclear factor kappa-B
OAT: Organic anion transporter
OATP: Organic anion transporting polypeptide
OCT: Organic cation transporter
OCTN: Organic cation/carnitine transporter
PBPK: Physiologically based pharmacokinetic
PCR: Polymerase chain reaction
PDTC: Pyrrolidinedithiocarbamate ammonium
PEPT2: Peptide transporter 2
P-gp: P-glycoprotein
PES: Polyethersulfone
PPARα: Peroxisome proliferator-activated receptor alpha
PTECs: Proximal tubular epithelial cells
PXR: Pregnane X receptor
qPCR: Quantitative PCR
RT-qPCR: Reverse transcription quantitative PCR
RXRα: Retinoid X receptor alpha
sIL-6Rα: Soluble interleukin-6 receptor alpha
SGLT2: Sodium-glucose cotransporter 2
SOCS3: Suppressor of cytokine signaling 3
TAB: Transforming growth factor-β–activated kinase 1 binding proteins
TAK1: Transforming growth factor-β–activated kinase 1
TNF-α: Tumor necrosis factor alpha
TRAF6: TNF receptor–associated factor 6
TYK2: Tyrosine kinase 2
UGT: UDP-glucuronosyltransferase
URAT1: Urate transporter 1

