## Supplemental File for "Regulation of Renal Transporters by Pro-inflammatory Cytokines in Human Proximal Tubular Epithelial Cells: Identification of the Perpetrator and Mechanisms"

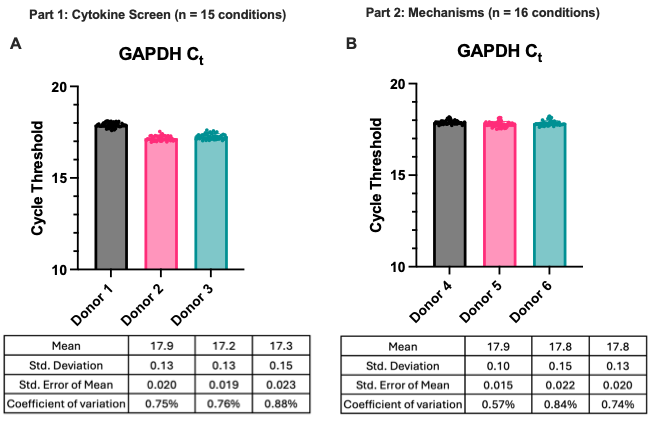

**Supplemental** Figure S1. GAPDH stability across donors and conditions. GAPDH cycle threshold (Ct) values in primary human PTECs remained stable within donors across treatments in both part 1 (A) and part 2 (B) of this study at the same amount of cDNA input (Donor 1, 4–6: 90 ng; Donor 2 and 3: 50 ng). Therefore, GAPDH was used for mRNA normalization throughout this study. Each point denotes average C_t_ values of technical triplicates. Bars are mean ± SD.

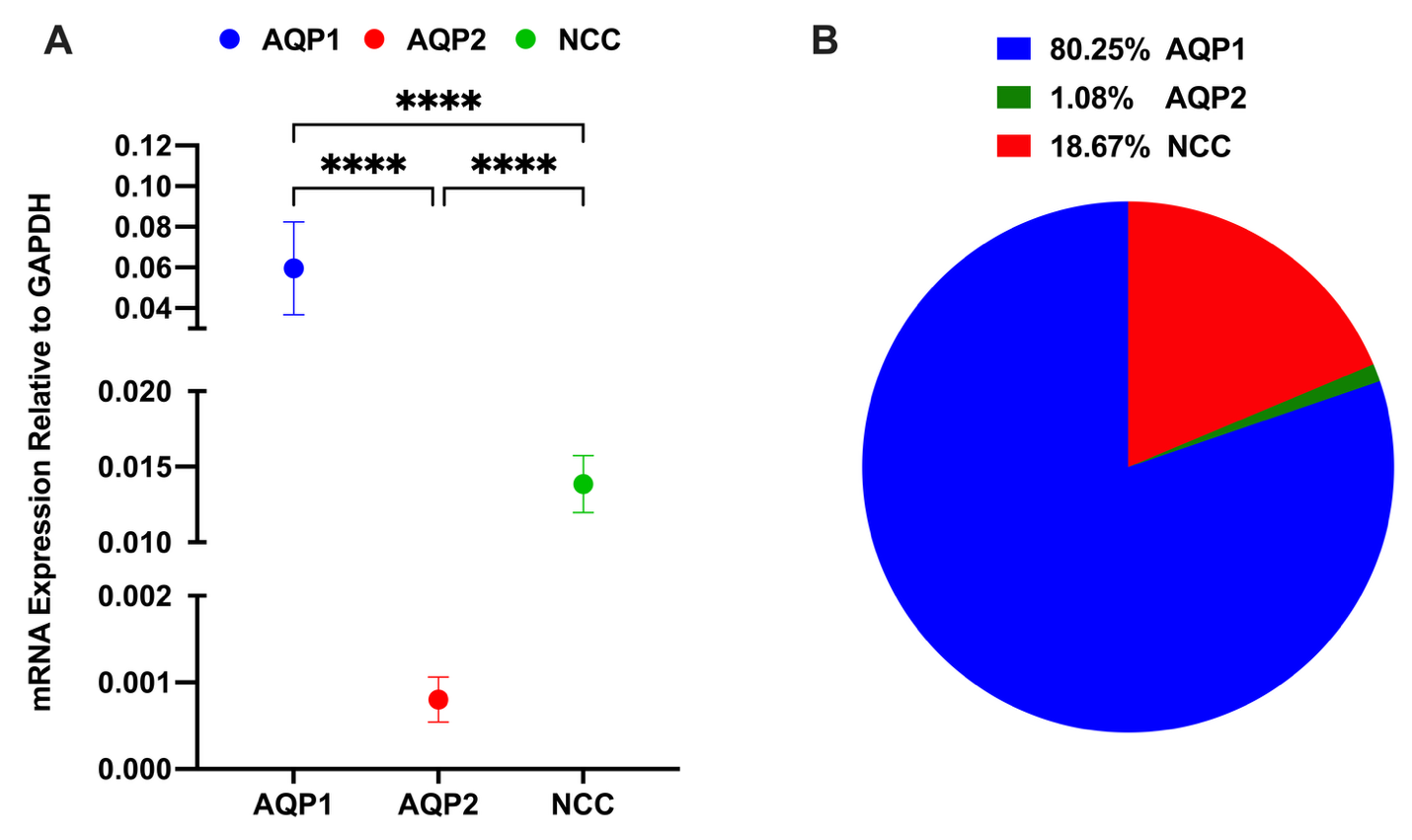

**Supplemental** Figure S2. Relative mRNA expression of renal cortical cell markers. Relative mRNA expression of three segment-specific markers (AQP1 for proximal tubule, AQP2 for cortical collecting duct, and NCC for distal tubule) in freshly isolated primary human PTECs at baseline shows a predominantly proximal phenotype (AQP1≫AQP2>NCC). (A) Data are means ± S.D. of each gene in vehicle-treated PTECs. Statistical significance were assessed using one-way ANOVA with Tukey’s multiple comparisons correction; (B) Pie chart summarizes proportional contribution of each marker.

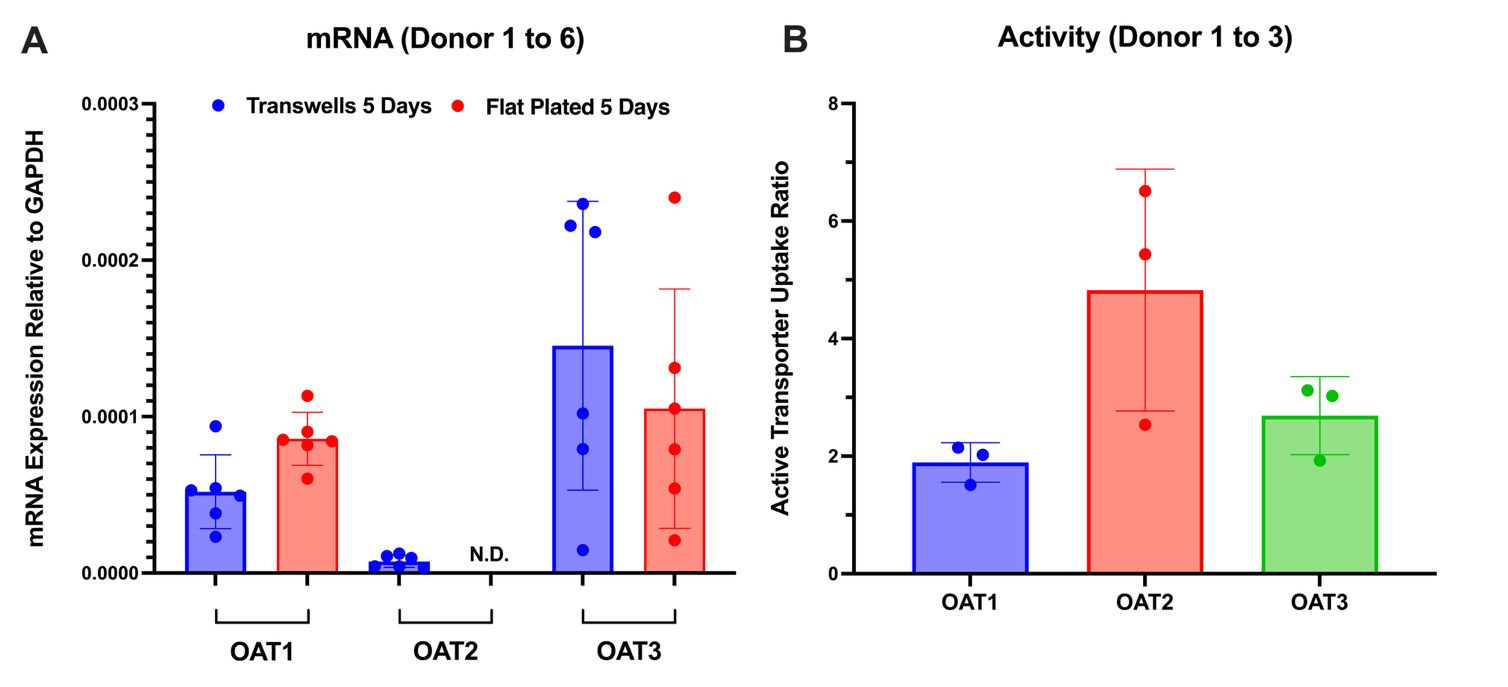

**Supplemental** Figure S3. OAT mRNA expression and activity in primary human PTECs cultured on Transwell inserts versus flat-plates. (A) Relative mRNA expression of OAT1/2/3 in vehicle-treated PTECs cultured on Transwells and flat-plates for five days. (B) Activity of OAT1/2/3 in vehicle-treated PTECs cultured on Transwells for five days (expressed as a ratio of substrate uptake without inhibitor to that with inhibitor; selective substrates and substrate-inhibitor pairs are shown in **Supplemental Table S3**). Data are mean ± SD (points denote donor, each tested in triplicate).

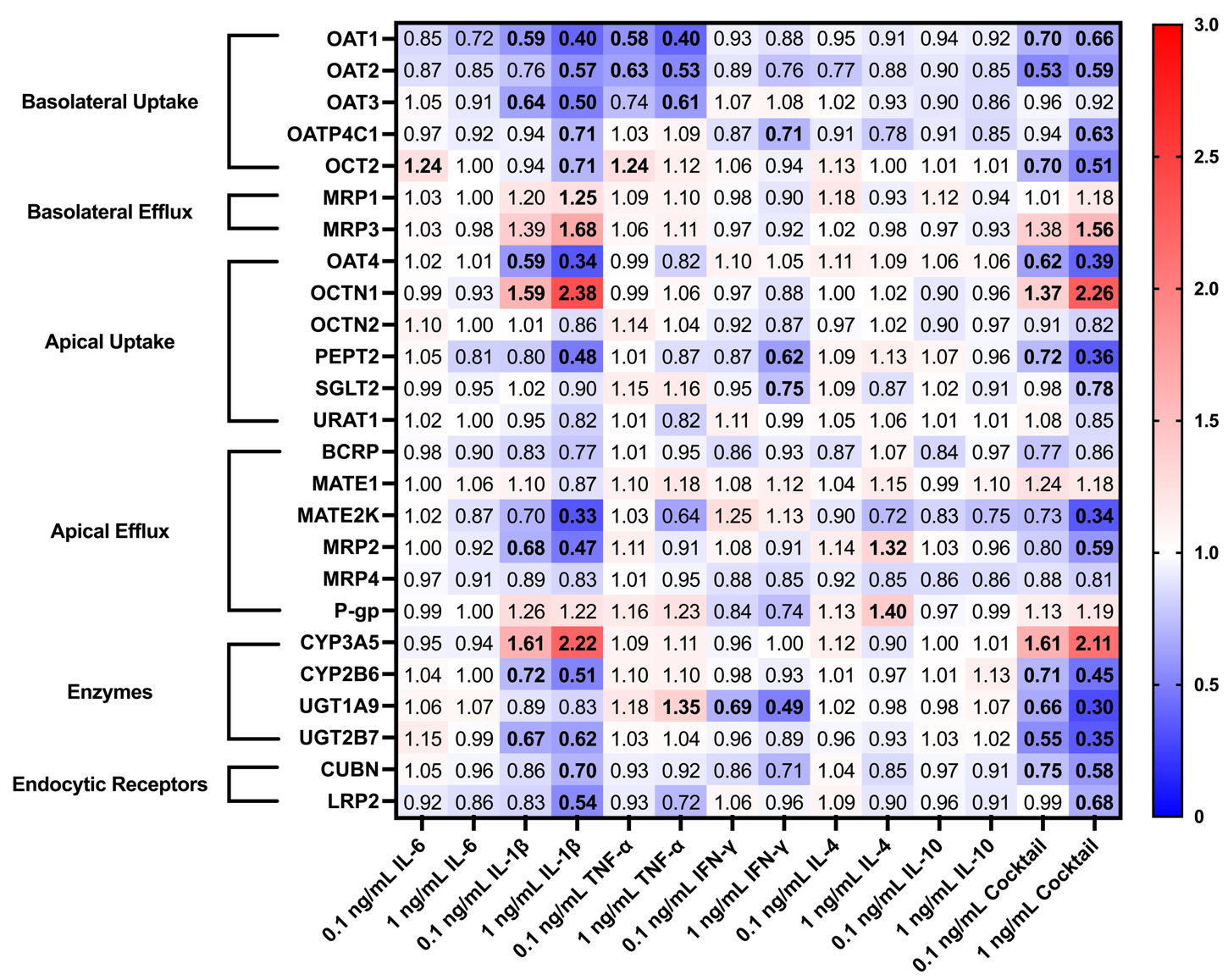

**Supplemental** Figure S4. Full heatmap summary of cytokine effects on the mRNA expression of renal drug transporters, drug metabolizing enzymes, and endocytic receptors. PTECs on Transwells were exposed to individual cytokines (IL-6, IL-1β, TNF-α, IFN-γ, IL-4, and IL-10) or the cytokine cocktail at 0.1 and 1 ng/mL in both chambers for 48 h (media replaced every 24 h). Heatmap shows average GAPDH-normalized mRNA expression relative to vehicle control (0.1% DPBS) across three donors. Gene expression significantly altered are bolded (*p<0.05, two-way ANOVA with Dunnett’s multiple comparison correction).

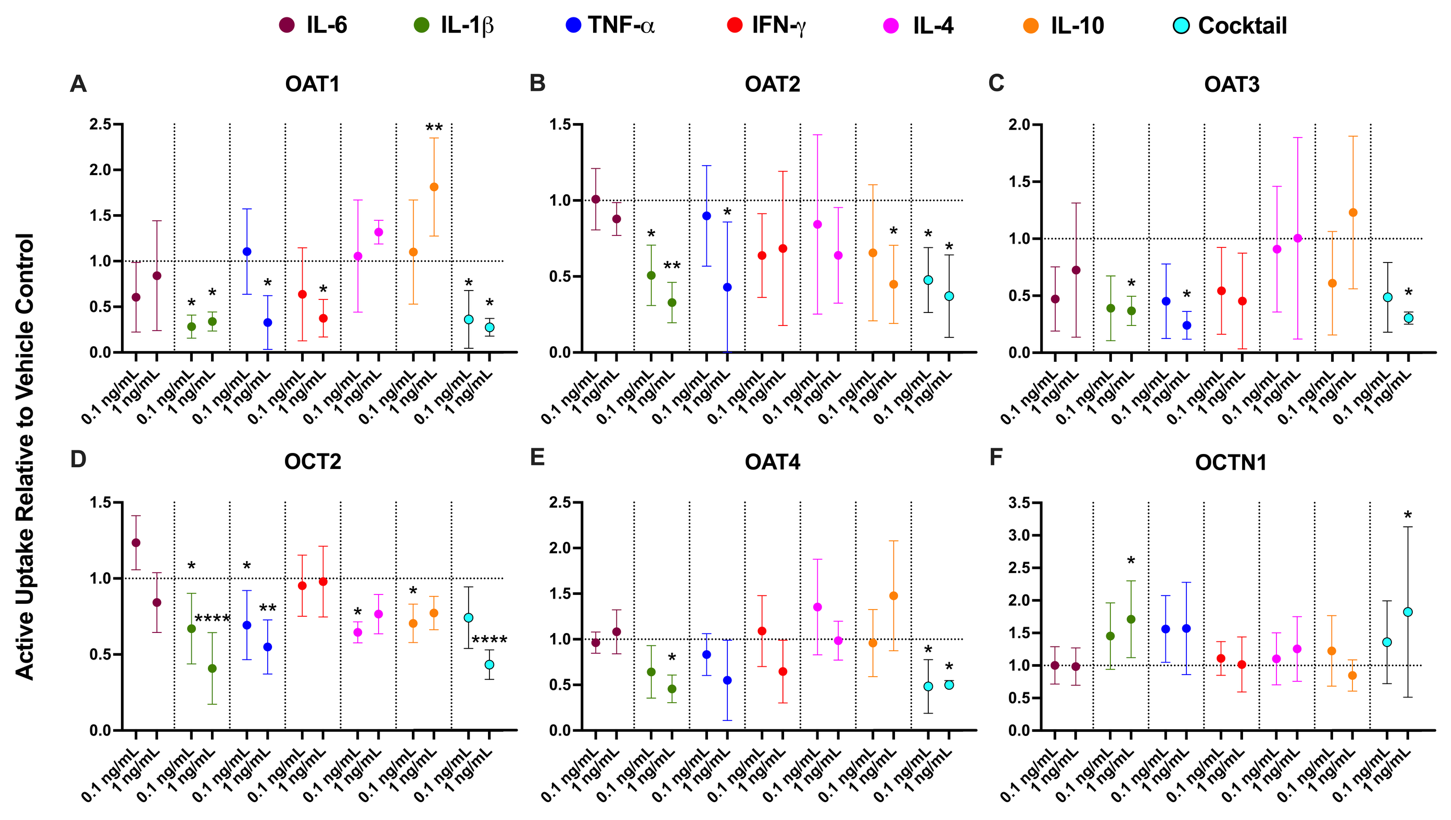

**Supplemental** Figure S5. **Renal uptake transporter activity in PTECs was modulated mainly by IL-1β and TNF-α.** PTECs on Transwells were exposed to individual cytokines (IL-6, IL-1β, TNF-α, IFN-γ, IL-4, and IL-10) or the cytokine cocktail at 0.1 and 1 ng/mL in both chambers for 48 h (media replaced every 24 h). Activity of uptake transporters ([**A**] OAT1, [**B**] OAT2, [**C**] OAT3, [**D**] OCT2, [**E**] OAT4, [**F**] OCTN1) is presented as the fraction of active uptake (determined by normalizing transporter-selective substrate uptake in the absence of inhibitors to that measured in the presence of inhibitors [**Supplemental Table S3**]) relative to vehicle-treated controls (0.1% DPBS; horizontal dashed line at y = 1). Similar to the mRNA results, IL-1β and TNF-α produced the broadest and largest changes in renal uptake transporter activity. Effects of the cytokine cocktail generally mirrored those of IL-1β. Data are mean ± SD from three donors (each quantified in triplicate). Statistical significance was assessed using two-way ANOVA with Dunnett’s multiple comparisons.

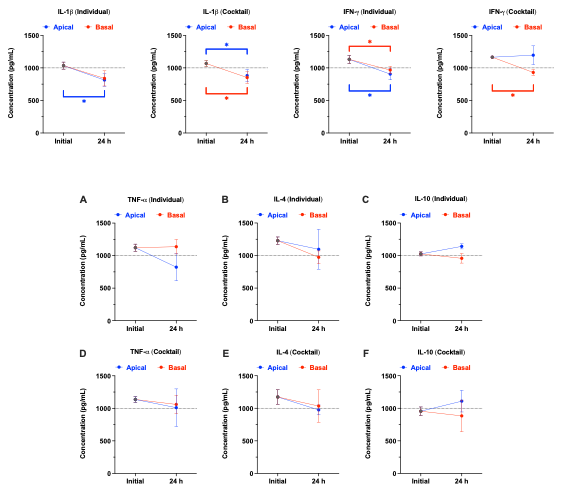

**Supplemental** Figure S6. Cytokine stability in apical and basal media over the first 24 h of exposure to PTECs on Transwells (ELISA). Primary human PTECs were treated as in Figure 4. Cytokines were added at 1 ng/mL to both chambers, and media were sampled at the start of treatment (“initial,” immediately after cytokine addition) and after 24 h (the first 24 h of a 48 h exposure period; media replaced at 24 h). Concentrations in the apical (blue) and basal (red) chambers were quantified by ELISA. Panels show concentrations of individual cytokines ([A] TNF-α, [B] IL-4, [C] IL-10) and of the same cytokines when included in the cocktail containing all six cytokines at 1 ng/mL each ([D] TNF-α, [E] IL-4, [F] IL-10). Data are ± SD from three donors (denoted by points, each tested in triplicate). Dotted lines denote the initial nominal 1 ng/mL concentration. Statistical significance was assessed using one-way ANOVA with Dunnett’s multiple comparisons correction versus the initial sample within each chamber. No significant decrease in concentration of TNF-α, IL-4, or IL-10 was detected over 24 h under either individual or cocktail conditions (p > 0.05). For other cytokines that showed statistically significant but modest decrease in concentration (IL-1β, IFN-γ), see **Figure 4B–E**.

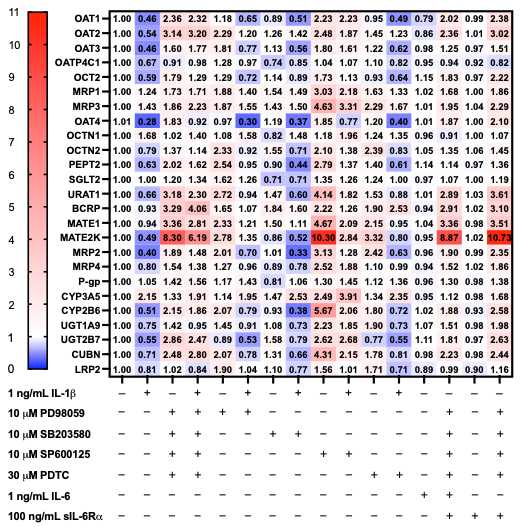

**Supplemental** Figure S7. Full heatmap summary of the regulation of renal DMET and endocytic receptor mRNA expression by IL-1β ± signaling pathway inhibitors and IL-6 trans-signaling with sIL-6Rα (vehicle-normalized). Heatmap shows average GAPDH-normalized mRNA expression relative to vehicle-treated controls (0.1% DPBS) across three donors. For IL-1β mechanism studies, primary human PTECs were treated for 48 h with 1 ng/mL IL-1β in both chambers of Transwells, with media replaced every 24 h. Cells were co-treated with small-molecule inhibitors targeting MEK/ERK (PD98059, 10 μM), p38 (SB203580, 10 μM), JNK (SP600125, 10 μM), or NF-κB (PDTC, 30 μM), either individually or as a four-inhibitor cocktail (“all +”). Inhibitor-only conditions were also included. For IL-6 trans-signaling studies, cells were treated for 48 h (media replaced every 24 h) with (i) 1 ng/mL IL-6; (ii) 1 ng/mL IL-6 + 100 ng/mL sIL‑6Rα + the inhibitor cocktail; (iii) 100 ng/mL sIL‑6Rα alone; and (iv) 100 ng/mL sIL‑6Rα + the inhibitor cocktail. IL-6 and the inhibitor cocktail were added to both apical and basal chamber of Transwells, while sIL-6Rα was only added to the basal chamber of Transwells to mimic its minimal *in vivo* renal filtration.

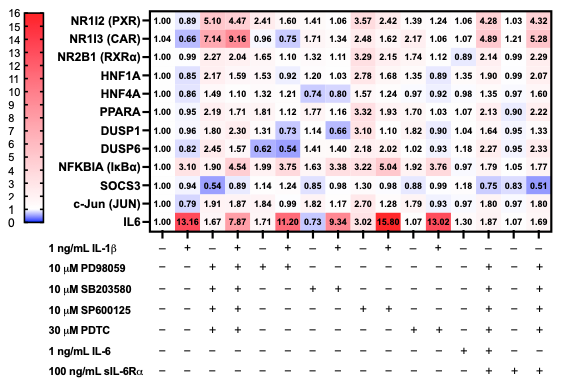

**Supplemental** Figure S8. GAPDH-normalized mRNA expression of nuclear receptors and pathway-engagement markers in PTECs after treatments with IL-1β ± pathway inhibitors or with IL-6 classic/trans-signaling related conditions, relative to vehicle control. Treatment conditions are detailed in **Supplemental** Figure S7. Heatmap shows averaged relative mRNA expression to vehicle (0.1% DPBS) across three donors. Nuclear receptors and transcriptional regulators (PXR, CAR, RXRα, HNF1α, HNF4α, PPARα) were included to assess secondary shifts in the regulatory pathway. IL-1β robustly induced IL6 and IκBα mRNA (≈ 13.2× and 3.1×, respectively), while downregulating mRNA of CAR mRNA (≈ 0.66×), PXR (≈ 0.89×), and DUSP6 (≈ 0.82×). The inhibitor cocktail strongly de-suppressed nuclear receptor mRNA (e.g., PXR ≈ 5×, CAR ≈ 7×), with modest upregulation of RXRα/HNF1A/HNF4A/PPARA mRNA and downregulation of SOCS3 mRNA (≈ 0.54×). IL-1β + cocktail restored CAR and PXR mRNA to above vehicle (CAR ≈ 9.2×; PXR ≈ 4.5×) despite persistent mRNA upregulation of IL-1β-responsive markers (IL6 ≈ 7.9×; IκBα ≈ 4.5×). PD98059 (MEK/ERK) downregulated DUSP6 mRNA (≈ 0.62× alone; ≈ 0.54× with IL-1β), consistent with ERK blockade. SB203580 (p38) downregulated DUSP1 mRNA under 1 ng/mL of IL-1β (≈ 0.66×), consistent with p38 inhibition. SP600125 (JNK) upregulated c-Jun (JUN) mRNA to above vehicle in the absence of IL-1β. In the presence of IL-1β, c-Jun mRNA returned to near-baseline (≈ 1.28×). IκBα mRNA remained upregulated with IL-1β in the presence of PDTC, indicating that IL-1β-driven IκBα mRNA upregulation was not blunted at this time point. Exogenous IL-6 alone produced minimal changes in SOCS3 mRNA on average (≈ 1.18×) and did not recapitulate the effects of IL-1β, consistent with limited IL-6 classic signaling in PTECs. IL-6 in the presence of sIL-6Rα showed relief of SOCS3 repression, consistent with IL-6 trans-signaling via STAT3 and suggests that JAK/STAT contributes minimally to renal transporter regulation (**Figures 7** and **Supplemental** Figure **S9)**. Data shown here represent a snapshot after feedback mechanisms have occurred, and are complementary to the functional IL-6 secretion data shown in **Figures 4, 6,** and **8**.

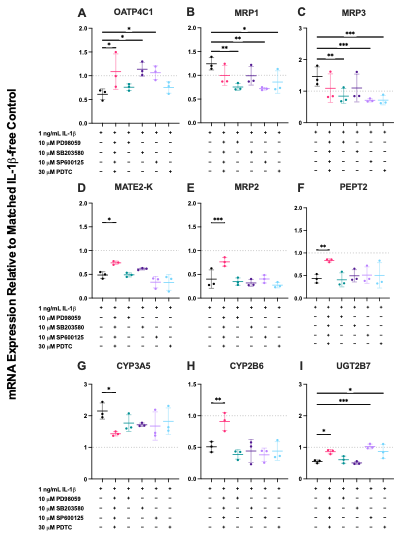

**Supplemental** Figure S9. Effect of pathway inhibitors on IL-1β–mediated regulation of renal DMET mRNA (matched-inhibitor normalized). Treatment conditions are detailed in **Supplemental** Figure **S7**. Here, the same dataset was normalized to the respective inhibitor baselines to emphasize rescue-of-effect. Statistical significance (p≤0.05, *p<0.01, **p<0.001, ***p<0.0001) was assessed using repeated-measures one-way ANOVA with Dunnett’s multiple comparisons.

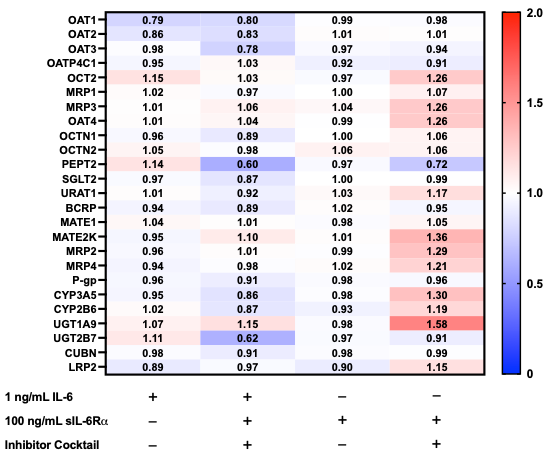

**Supplemental** Figure S10. Full heatmap summary of the effect of IL-6 classic and trans-signaling on renal DMET and endocytic receptor mRNA (matched-inhibitor–normalized). Treatment conditions are detailed in **Supplemental Figure S7**. Heatmap shows averaged GAPDH-normalized mRNA data across three donors (each quantified in technical triplicate) normalized to the respective inhibitor baselines to emphasize rescue-of-effect. Both classic and trans-signaling of IL-6 did not significantly affect the mRNA expression of renal DMETs and endocytic receptors, except for PEPT2 and UGT2B7 (**Figure 7**).

### Supplemental Table S1. Chemicals and Reagents.

| **Category** | **Reagent** | **Supplier** |
| --- | --- | --- |
| Media and buffers | Dulbecco’s Modified Eagle’s Medium/Nutrient Mixture F-12 (DMEM/F-12) powder without glucose | United States Biological (Salem, MA, USA) |
|  | D-(+)-Glucose | Sigma-Aldrich (St. Louis, MO, USA) |
|  | HEPES |  |
|  | Sodium bicarbonate (NaHCO_3_) |  |
|  | Sodium hydroxide (NaOH) |  |
|  | HBSS with Ca2+ and Mg2+ (HBSS^+/+^) | Thermo Fisher Scientific (Waltham, MA, USA) |
|  | HBSS without Ca2+ and Mg2+ (HBSS^–/–^) |  |
| Enzymes and dissociation reagents | Collagenase type IV (powder) | Thermo Fisher Scientific (Waltham, MA, USA) |
|  | Dispase II (powder) |  |
|  | Calcium chloride (CaCl_2_) | Sigma-Aldrich (St. Louis, MO, USA) |
|  | Ethylenediaminetetraacetic acid disodium salt dihydrate (EDTA) disodium salt dihydrate |  |
| Supplements and additives | Antibiotic–Antimycotic (100×) | Thermo Fisher Scientific (Waltham, MA, USA) |
|  | Bovine serum albumin (BSA) |  |
|  | Insulin-Transferrin-Selenium (ITS-G, 100×) |  |
|  | Matrigel Growth Factor Reduced (Phenol Red-free) | Corning (Corning, NY, USA) |
|  | HumanKine recombinant human EGF | Proteintech (Rosemont, IL, USA) |
|  | Hydrocortisone (Cortisol) | Sigma-Aldrich (St. Louis, MO, USA) |
|  | A83-01 | MedChemExpress (Monmouth Junction, NJ, USA) |
|  | Y-27632 |  |
|  | Triiodothyronine (T3) |  |
| Solvents and LC–MS reagents | Acetonitrile (LC–MS grade) | Thermo Fisher Scientific (Waltham, MA, USA) |
|  | Dimethyl sulfoxide (DMSO) |  |
|  | Formic acid (LC–MS grade) |  |
| Cultureware and plasticware | 96-well PCR plates | Thermo Fisher Scientific (Waltham, MA, USA) |
|  | Nalgene Rapid-Flow Sterile Disposable Bottle Top Filters with 0.2 μm PES Membrane |  |
|  | Falcon 70 μm cell strainers | Corning (Corning, NY, USA) |
|  | Transwell-Clear Inserts, Polyester (PET) membrane |  |
| Density media | Percoll density gradient medium | Cytiva (Marlborough, MA, USA) |
| Cytokines and cytokine receptors | HumanKine recombinant IL-6 | Proteintech (Rosemont, IL, USA) |
|  | HumanKine recombinant IL-1β |  |
|  | HumanKine recombinant TNF-α |  |
|  | HumanKine recombinant IFN-γ |  |
|  | HumanKine recombinant IL-4 |  |
|  | HumanKine recombinant IL-10 |  |
|  | Recombinant Human IL-6R alpha Protein | R&D Systems (Minneapolis, MN, USA) |
| RNA isolation, cDNA Synthesis, and RT-qPCR | Dithiothreitol (DTT) | MedChemExpress (Monmouth Junction, NJ, USA) |
|  | 96-well PCR and assay microplates | Thermo Fisher Scientific (Waltham, MA, USA) |
|  | PureLink RNA Mini Kit |  |
|  | PureLink DNase Set |  |
|  | High-Capacity cDNA Reverse Transcription Kit |  |
|  | TaqMan Fast Advanced Master Mix |  |
|  | TaqMan Gene Expression Assays (Assay IDs in **Supplemental Table S3**) |  |
|  | DNase I |  |
| Transporter probes (non-radioactive) | Glycochenodeoxycholic acid-sulfate (GCDCA-S) | LGC Group (Teddington, UK) |
|  | Levocetirizine hydrochloride | MedChemExpress (Monmouth Junction, NJ, USA) |
| Radioactive transporter probes and scintillation | [^3^H]nicotinic acid (50 Ci/mmol, 20 mM) | American Radiolabeled Chemicals (St. Louis, MO, USA) |
|  | [^3^H]cidofovir (21.7 Ci/mmol, 46.1 mM) | Moravek (Brea, CA, USA) |
|  | [^3^H]atenolol (3.3 Ci/mmol, 0.303 mM) |  |
|  | [^3^H]ergothioneine (0.4 Ci/mmol, 2.5 mM) |  |
|  | Ecoscint ORIGINAL | National Diagnostics (Atlanta, GA, USA) |
| Transporter inhibitors and small molecules | Bromosulfophthalein disodium salt (BSP) | MedChemExpress (Monmouth Junction, NJ, USA) |
|  | Probenecid |  |
|  | Pyrimethamine |  |
|  | Cyclosporine A |  |
|  | Mitoxantrone |  |
|  | Quercetin |  |
|  | Ergothioneine |  |
|  | Fexofenadine hydrochloride |  |
|  | PD98059 |  |
|  | SB203580 |  |
|  | SP600125 |  |
|  | Pyrrolidinedithiocarbamate ammonium (PDTC) |  |
| Protein quantification | Pierce Bicinchoninic Acid (BCA) Protein Assay Kit | Thermo Fisher Scientific (Waltham, MA, USA) |
| ELISA kits | Human IL-6 ELISA Kit | Proteintech (Rosemont, IL, USA) |
|  | Human IL-1β ELISA Kit |  |
|  | Human TNF-α ELISA Kit |  |
|  | Human IFN-γ ELISA Kit |  |
|  | Human IL-4 ELISA Kit |  |
|  | Human IL-10 ELISA Kit |  |
|  | Human IL-6R alpha ELISA Kit | R&D Systems (Minneapolis, MN, USA) |

### Supplemental Table S2. PTEC donor demographics. Donor kidneys were disease-free. Kidneys were deemed unsuitable for clinical transplantation for allocation or quality reasons unrelated to intrinsic kidney disease (e.g., not allocated within the clinical time window, prolonged cold ischemia, or donor serologies incompatible with available recipients).

| **Donor** | **Age** | **BMI (kg/m^2^)** | **Sex** | **Race** | **Cause of death** | **Alcohol use*** | **Tobacco use*** | **Substance use*** |
| --- | --- | --- | --- | --- | --- | --- | --- | --- |
| Donor 1^a^ | 43 | 40.9 | Female | White | Anoxia | No | No | No |
| Donor 2^a^ | 69 | 20.4 | Male | White | Head Trauma | Yes | Yes | No |
| Donor 3^a^ | 49 | 25.5 | Female | White | Anoxia | No | No | No |
| Donor 4^b^ | 39 | 31.2 | Female | White | Anoxia | No | No | No |
| Donor 5^b^ | 58 | 24.5 | Female | White | Respiratory | No | Yes | No |
| Donor 6^b^ | 49 | 27.8 | Male | White | Stroke | No | No | No |

*Current or recent use as recorded in Organ Procurement Organization donor charts.

^a^ Used for Part 1 of this study.

^b^ Used for Part 2 of this study.

**Supplemental** Table S3. Substrates and inhibitors used for uptake assays in primary human PTECs. Selective substrates and substrate-inhibitor pairs used to measure individual uptake transporter activity in PTECs are listed. Adapted from Tsang et al. (1).

| **Target Uptake Transporter** | **Selective Substrate or**  **Substrate-Inhibitor* Pair** | **Inhibitor of the Targeted Uptake Transporter** |
| --- | --- | --- |
| OAT1 | 46.1 nM [^3^H]cidofovir | >200 $\mu$M probenecid |
| OAT2 | 20 nM [^3^H]nicotinic acid (substrate)  + 25 $\mu$M quercetin (inhibitor) | >200 $\mu$M BSP |
| OAT3 | 5 μM GCDCA-S (substrate)  + 50 $\mu$M cyclosporine A (inhibitor) | >200 $\mu$M probenecid |
| OAT4 | 1 μM levocetirizine | >200 $\mu$M BSP |
| OCT2 | 303 nM [^3^H]atenolol (substrate)  + 25 $\mu$M mitoxantrone (inhibitor) | >200 $\mu$M pyrimethamine |
| OCTN1 | 2.5 μM [^3^H]ergothioneine | 1 mM ergothioneine |

*****For substrate-inhibitor pairs, the inhibitor blocks non-target transporters involved in the flux of the substrate in PTECs, allowing selective measurement of the indicated uptake transporter.

**Supplemental Table S4. TaqMan Assay IDs**

| **Gene** | **Assay ID** |
| --- | --- |
| OAT1 (*SLC22A6*) | Hs00537914_m1 |
| OAT2 (*SLC22A7*) | Hs00198527_m1 |
| OAT3 (*SLC22A8*) | Hs01056646_m1 |
| OATP4C1 (*SLCO4C1*) | Hs00698884_m1 |
| OCT2 (*SLC22A2*) | Hs01010726_m1 |
| MRP1 (*ABCC1*) | Hs01561483_m1 |
| MRP3 (*ABCC3*) | Hs00978452_m1 |
| OAT4 (*SLC22A11*) | Hs00945829_m1 |
| OCTN1 (*SLC22A4*) | Hs00268200_m1 |
| OCTN2 (*SLC22A5*) | Hs00929869_m1 |
| PEPT2 (*SLC15A2*) | Hs01113665_m1 |
| SGLT2 (*SLC5A2*) | Hs00894642_m1 |
| URAT1 (*SLC22A12*) | Hs01030727_m1 |
| BCRP (*ABCG2*) | Hs01053790_m1 |
| MATE1 (*SLC47A1*) | Hs00217320_m1 |
| MATE2-K (*SLC47A2*) | Hs00945652_m1 |
| MRP2 (*ABCC2*) | Hs00960489_m1 |
| MRP4 (*ABCC4*) | Hs00988721_m1 |
| P-gp (*ABCB1*) | Hs00184500_m1 |
| CYP3A5 (*CYP3A5*) | Hs00241417_m1 |
| CYP2B6 (*CYP2B6*) | Hs03044634_m1 |
| UGT1A9 (*UGT1A9*) | Hs02516855_sH |
| UGT2B7 (*UGT2B7*) | Hs00426592_m1 |
| Cubilin (*CUBN*) | Hs00153607_m1 |
| Megalin (*LRP2*) | Hs00189742_m1 |
| PXR (*NR1I2*) | Hs01114267_m1 |
| CAR (*NR1I3*) | Hs00901571_m1 |
| RXRα (*NR2B1*) | Hs01067640_m1 |
| HNF1α (*HNF1A*) | Hs00167041_m1 |
| HNF4α (*HNF4A*) | Hs00230853_m1 |
| PPARα (*PPARA*) | Hs00947536_m1 |
| DUSP1 (*DUSP1*) | Hs00610256_g1 |
| DUSP6 (*DUSP6*) | Hs00169257_m1 |
| IκBα (*NFKBIA*) | Hs00153283_m1 |
| SOCS3 (*SOCS3*) | Hs02330328_s1 |
| c-Jun (*JUN*) | Hs01103582_s1 |
| IL-6 (*IL6*) | Hs00174131_m1 |
| GAPDH (*GAPDH*) | Hs99999905_m1 |

### Supplemental Table S5. Baseline GAPDH-normalized mRNA expression of renal transporters in vehicle-treated PTECs on Transwells after 5 days in culture.

|  |  | **Part 1** | | | **Part 2** | | |
| --- | --- | --- | --- | --- | --- | --- | --- |
|  |  | Donor 1 | Donor 2 | Donor 3 | Donor 4 | Donor 5 | Donor 6 |
| **Apical Transporters** | **OAT1** | 9.38E-05 | 2.32E-05 | 3.81E-05 | 5.43E-05 | 4.95E-05 | 5.29E-05 |
|  | **OAT2** | 1.08E-05 | 1.24E-05 | 3.48E-06 | 4.24E-06 | 4.28E-06 | 9.68E-06 |
|  | **OAT3** | 2.18E-04 | 1.47E-05 | 7.93E-05 | 2.22E-04 | 2.36E-04 | 1.02E-04 |
|  | **OATP4C1** | 1.22E-02 | 1.54E-02 | 9.85E-03 | 2.89E-02 | 2.39E-02 | 1.11E-02 |
|  | **OCT2** | 5.53E-03 | 1.08E-02 | 4.78E-03 | 6.79E-03 | 7.33E-03 | 6.94E-03 |
|  | **MRP1** | 4.66E-03 | 1.33E-02 | 7.57E-03 | 9.68E-03 | 9.66E-03 | 1.62E-02 |
|  | **MRP3** | 7.70E-03 | 2.35E-02 | 4.51E-03 | 1.19E-02 | 1.29E-02 | 1.01E-02 |
| **Basolateral Transporters** | **OAT4** | 1.48E-03 | 5.08E-04 | 2.21E-04 | 3.17E-03 | 3.74E-03 | 1.34E-03 |
|  | **OCTN1** | 4.03E-04 | 8.55E-04 | 4.19E-04 | 9.37E-04 | 7.85E-04 | 4.87E-04 |
|  | **OCTN2** | 3.48E-03 | 3.32E-03 | 1.93E-03 | 5.59E-03 | 6.31E-03 | 3.93E-03 |
|  | **PEPT2** | 1.14E-04 | 3.24E-04 | 1.75E-04 | 1.63E-04 | 1.64E-04 | 3.05E-04 |
|  | **SGLT2** | 1.46E-04 | 2.09E-04 | 6.27E-05 | 2.63E-04 | 2.37E-04 | 1.00E-04 |
|  | **URAT1** | 2.63E-04 | 5.98E-05 | 6.63E-05 | 1.87E-04 | 1.98E-04 | 1.46E-04 |
|  | **BCRP** | 9.40E-05 | 1.04E-04 | 4.31E-05 | 6.31E-05 | 7.50E-05 | 4.16E-05 |
|  | **MATE1** | 5.32E-04 | 2.92E-04 | 2.33E-04 | 1.47E-03 | 1.52E-03 | 8.65E-04 |
|  | **MATE2K** | 9.02E-04 | 5.58E-05 | 1.58E-04 | 9.78E-04 | 8.90E-04 | 2.43E-04 |
|  | **MRP2** | 1.70E-03 | 1.01E-04 | 2.27E-04 | 1.74E-03 | 1.88E-03 | 1.18E-03 |
|  | **MRP4** | 6.48E-03 | 6.62E-03 | 5.01E-03 | 5.95E-03 | 6.57E-03 | 5.36E-03 |
|  | **P-gp** | 6.55E-03 | 4.82E-03 | 4.23E-03 | 1.64E-02 | 1.65E-02 | 9.19E-03 |
